# Music exposure reduces anxiety- and depression-like behavior in rodents: a systematic review and multilevel meta-analysis

**DOI:** 10.64898/2026.02.27.708573

**Authors:** Santiago Ortega, Anna Lenz, Erick Lundgren, Ayumi Mizuno, Sergio Poo Hernandez, Shinichi Nakagawa, Malgorzata Lagisz

## Abstract

Anxiety and depressive disorders impose a major global burden, prompting interest in non-pharmacological interventions that may influence affective processes. Music exposure has often been reported to affect anxiety- and depression-like behaviors, but preclinical findings remain heterogeneous and have not been quantitatively synthesized. Prior work has also focused almost entirely on mean behavioral responses, largely overlooking inter-individual variability as a biologically meaningful dimension. We conducted a preregistered systematic review and multilevel meta-analysis of experimental studies testing music exposure in laboratory rodents. Following PRISMA and PRISMA-EcoEvo guidelines, we synthesized 298 effect sizes from 20 studies using multilevel models to account for non-independence among effect sizes. We quantified effects on mean behavior with the log response ratio (ln*RR*) and effects on variability with the log variability ratio (ln *V R*). Overall, music exposure was associated with a statistically significant reduction in anxiety- and depression-like behaviors, corresponding to an average decrease of about 18% relative to controls. This mean effect was detected across outcome types and life stages despite substantial heterogeneity. By contrast, music exposure did not produce a statistically significant overall change in inter-individual behavioral variability. Instead, variability responses were context dependent: behavioral assay type and music meta-genre significantly moderated ln *V R*, with anxiety-like assays tending to show increased variability and depression-like assays tending to show reduced variability under music exposure. These results suggest that music exposure reliably shifts average affect-related behavior without uniformly changing behavioral stability across individuals. Because the evidence comes mainly from short-term exposures in young adult laboratory rodents, generalization beyond similar contexts should remain cautious.

## 1 Introduction

Anxiety and depressive disorders are among the most prevalent and disabling psychiatric conditions worldwide. Anxiety disorders are characterized by persistent fear, hyper-vigilance, and heightened stress reactivity, whereas depressive disorders involve sustained low mood, anhedonia, and impaired motivation. Together, these conditions affect hundreds of millions of individuals globally and are major contributors to reduced quality of life and economic burden (Airaksinen et al., 2024; Ferrari, 2022; Ridley et al., 2020). Their high prevalence and chronic nature have spurred extensive efforts to understand the neural mechanisms underlying affective dysregulation and to develop effective treatments.

Pharmacological therapies remain a cornerstone of treatment, but their efficacy is variable and often accompanied by side effects or incomplete symptom relief (Boulenger, 2004; Kelly and Mezuk, 2017). As a result, there has been growing interest in complementary and non-pharmacological approaches that may modulate affective state through engagement of neural systems involved in emotion regulation and stress responsiveness. Among these, music exposure has received substantial attention (de Witte et al., 2022). In humans, music reliably engages limbic, paralimbic, and cortical networks implicated in emotion, reward, and stress regulation, and can elicit robust physiological and affective responses (Arnold et al., 2024; Koelsch, 2014; Levitin and Tirovolas, 2009). Clinical and experimental studies suggest that music can reduce self-reported anxiety and depressive symptoms, at least transiently, and may enhance emotional regulation across a range of contexts (Fancourt and Finn, 2019; Thoma et al., 2013).

Although the neural mechanisms through which music engages affective and stress-related systems are increasingly well characterized, translating these effects into reliable and generalizable therapeutic applications remains challenging (Fancourt et al., 2014; Koelsch, 2014). Individual differences in musical preference, prior experience, clinical condition, and cultural context introduce substantial variability in outcomes, complicating assessment of when music exposure can be used consistently to modulate anxiety- and depression-related processes (Balkwill and Thompson, 1999; McDermott et al., 2016). Preclinical animal models therefore provide a valuable complementary framework for addressing these limitations.

Rodent models allow precise control over auditory exposure, developmental timing, and environmental conditions, while enabling standardized assessment of anxiety- and depression-like behaviors. Behavioral assays such as Elevated Plus Maze, Open Field Test, Light–Dark Box, Forced Swim Test, Tail Suspension Test, and Sucrose Preference Test are widely used to probe stress reactivity, avoidance behavior, and coping strategies under controlled conditions (e.g., Cryan and Holmes, 2005; Slattery and Cryan, 2012). These models are central to behavioral neuroscience because they permit systematic investigation of how environmental inputs interact with neural and endocrine systems implicated in affective regulation, including stress-related and motivational pathways (Mcewen et al., 2007; Nestler et al., 2002).

Within this experimental framework, a growing literature has examined the effects of music exposure on rodent behavior. Many studies report reductions in anxiety- or depression-like behaviors following music exposure, consistent with the idea that structured auditory environments can modulate stress-related processes (e.g., Chikahisa et al., 2007; Escribano et al., 2014). However, studies differ in the type of music used, exposure duration and timing, developmental stage at exposure, behavioral assays employed, control conditions, and whether behaviors are assessed under baseline conditions or following experimentally induced stress (Kühlmann et al., 2018; Le et al., 2025). Consequently, reported effects vary widely. As a result, it remains unclear whether music exposure exerts a generalizable effect on affect-related behavior or whether its influence is highly context dependent.

A further limitation of the existing literature is its near-exclusive focus on changes in mean behavioral responses. Yet behavioral variability is itself a biologically meaningful property (Westneat et al., 2015). Changes in variance can reflect altered stability, sensitivity to environmental perturbations, or heterogeneity in individual stress responsiveness. Environmental manipulations may reduce variability by stabilizing behavioral responses or increase variability by amplifying individual differences (Sih et al., 2015). Whether music exposure affects not only average behavior but also the consistency of behavioral responses has not been systematically examined.

Recent advances in meta-analytic methodology allow a more rigorous synthesis of this evidence. Multilevel meta-analytic models accommodate multiple, non-independent effect sizes from the same study while appropriately partitioning sources of variance (Nakagawa and Santos, 2012). Effect size metrics such as the log response ratio (ln *RR*) enable comparison across diverse behavioral measures, whereas variance-based metrics such as the log variability ratio (ln *V R*) permit explicit evaluation of changes in behavioral variability (Nakagawa et al., 2015; Senior et al., 2020). Together, these approaches allow assessment of both the average effects of music exposure on affect-related behavior and the consistency of those effects across individuals and experimental contexts.

Here, we present a preregistered systematic review and multilevel meta-analysis of experimental studies examining the effects of music exposure on anxiety- and depression-like behaviors in laboratory rodents. Reported in accordance with PRISMA and PRISMA-EcoEvo guidelines (O’Dea et al., 2021; Page et al., 2021), this synthesis quantifies both mean behavioral effects and changes in behavioral variability, identifies sources of heterogeneity, and evaluates the robustness of reported effects. By clarifying the magnitude, consistency, and context dependence of music-induced behavioral changes under controlled conditions, this study provides a quantitative and conceptual framework for interpreting preclinical evidence relevant to affective regulation and, more broadly, to hypotheses about sensory modulation of anxiety- and depression-related processes.

## 2 Methods

### 2.1 Literature searching

We used the PECO (Population, Exposure, Comparator, Outcome; Table 1) framework (Foo et al., 2021; Rickard et al., 2005) to define the scope of the review and guide literature searching. We searched Web of Science, Scopus, PubMed, and OpenAlex. OpenAlex was included to improve coverage of newly indexed and non-traditionally published records that may not yet be fully integrated into subscription-based databases. Our search strings targeted experimental studies examining the effects of music exposure on anxiety- and depression-like behaviors in laboratory rodents. No temporal restrictions were applied to publication dates. We provide database-specific search strings in the Supplementary Material (S1). We evaluated search sensitivity by estimating relative recall against a benchmark set of 10 pre-identified experimental studies (Lagisz et al., 2025).

**Table 1:** Descriptions of the population, exposure, comparator, and outcome (PECO) used to define the scope of this study.

| **PECO** | **Description** |
| --- | --- |
| **Population** | Laboratory mice (*Mus musculus*) and rats (*Rattus norvegicus*). |
| **Exposure** | Exposure to music: structured auditory stimuli with organized temporal patterns (melody/rhythm). Excludes unstructured noise. |
| **Comparator** | Non-music control conditions (silence, white noise, or ambient sound). |
| **Outcome** | Anxiety-like and depression-like behaviors. |

To enhance coverage and identify gray literature, we additionally searched Google Scholar and the Bielefeld Academic Search Engine (BASE). Search strings were translated into Spanish, Japanese, Polish, and Russian, and Google Scholar screening was restricted to the first 50 results per language query. The search strategy, screening protocol, and planned analyses were preregistered on the Open Science Framework (https://osf.io/mv78t).

### 2.2 Eligibility criteria and screening

Study eligibility criteria followed the preregistered protocol and PECO (Population, Exposure, Comparator, Outcome; Table 1) framework. In brief, to be eligible for inclusion, studies had to report experiments assessing the effects of music exposure on anxiety- or depression-like behaviors in laboratory mice or rats. We excluded studies using wild-caught animals. For genetically modified models or animals subjected to invasive interventions (e.g., surgical or pharmacological procedures), we only included data from wild-type or sham-operated groups, if available; valid comparisons were restricted to music-exposed versus control animals within the same genetic or procedural background. Studies relying exclusively on physiological or histological outcomes without standardized behavioral assays were excluded. We also excluded studies lacking a concurrent control group because treatment effects could not be estimated without a comparator. We used Rayyan (Ouzzani et al., 2016) for screening titles and abstracts from bibliographic records. English-language records were screened independently by at least two reviewers (S.O., M.L., A.L., A.M., E.L., S.N., and S.P.), whereas non-English records were screened by a single reviewer with relevant language expertise. Full-text screening followed the same procedure, also in Rayyan. Screening of non-English language records and studies was conducted by reviewers with relevant language expertise (e.g., S.P., A.L., and S.O. for Spanish; M.L. for Polish and Russian; A.M. for Japanese).

### 2.3 Data collection

We extracted four categories of data from each eligible study. Data extraction was conducted by S.O., and a random 10% subset of studies (two studies) was independently cross-checked for accuracy against the original reports. We extracted: (i) Bibliographic information (title, authors, year, source, DOI). (ii) Music exposure and experimental conditions, including music type and, when reported, specific piece/composer; exposure duration (hours/day) and total exposure length (days); timing relative to behavioral testing (before, concurrent, or both); and light–dark phase during exposure; acoustic characteristics (sound level in dB; frequency in Hz) when available; control conditions (silence, ambient sound, or white noise) and their acoustic properties when reported; additional experimental manipulations (e.g., housing or stress procedures) when applied equivalently across treatment and control groups. (iii) We extracted numerical behavioral outcome data needed for effect size computation, including sample sizes, group means, and measures of dispersion (SD, SE, or confidence intervals). Outcomes were drawn from standardized behavioral assays only (e.g., Elevated Plus Maze, Open Field Test, Light–Dark Box, Forced Swim Test, Tail Suspension Test, Sucrose Preference Test). When numerical values were reported only graphically, we extracted data from plots using the *metaDigitise* R package (Pick et al., 2019). Outcomes were classified as reflecting innate behavior (e.g., approach–avoidance to exposed, well-lit spaces) or behavior induced by experimental procedures (e.g., chronic unpredictable mild stress) (Cryan and Holmes, 2005). (iv) Study population characteristics, including species, strain, sex, age at exposure/testing, randomization, blinding, sham procedures, attrition reporting, and completeness of outcome reporting. When information critical for effect size calculation was unclear— most commonly when sample sizes were reported as ranges rather than exact values—we contacted study authors by email (two studies) to request clarification. We received no responses to our emails, and thus we used the median of the reported sample-size range for effect size computation.

### 2.4 Deviations from the preregistered protocol

Three minor refinements were implemented during data extraction to accommodate heterogeneity in reporting practices. These refinements did not alter the PECO framework, eligibility criteria, outcome definitions, or primary analytical strategy and were implemented prior to data analysis. First, to model statistical dependence among effect sizes, we coded the comparison structure of each effect size as independent-group or dependent (within-subject/repeated-measures). This extended the preregistered metadata structure and enabled appropriate sampling-variance specification when multiple effect sizes were derived from the same experimental cohort.

Second, to harmonize heterogeneous reporting, we categorized exposure duration into acute, short-term, medium-term, and long-term classes, and grouped music stimuli into broad meta-genre categories based on historical context and dominant musical features (definitions in Table S1). These classifications were derived from reported information and used to facilitate interpretable moderator analyzes.

Third, we recorded whether key stimulus parameters (e.g., loudness, frequency, light–dark phase) were explicitly reported. Because loudness and frequency were frequently under-reported, we coded non-reporting explicitly rather than treated as missing to quantify reporting gaps.

### 2.5 Effect size calculation

We quantified the effects of music exposure using effect size metrics that capture both differences in mean behavioral responses and differences in behavioral variability between music-exposed and control groups.

#### 2.5.1 Log Response Ratio (ln *RR*)

Differences in mean behavioral responses were quantified using the natural logarithm of the response ratio (Hedges et al., 1999; Lajeunesse, 2011; Senior et al., 2020). To ensure consistent interpretation across assays, behavioral outcomes were coded so that negative ln *RR* values indicate a beneficial effect of music exposure (i.e., reduced anxiety- or depression-like behavior). For metrics where higher values reflect reduced anxiety/depression (e.g., time in open arms); the sign of ln *RR* was reversed prior to analysis.

To reduce small-sample bias arising from sampling error in group means, we used the bias-corrected estimator of the log response ratio. For each comparison, ln *RR* was calculated as

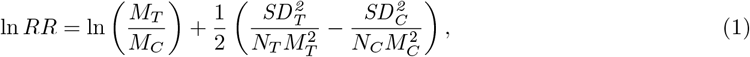

where *M*_*T*_ and *M*_*C*_ denote the sample means, *SD*_*T*_ and *SD*_*C*_ the sample standard deviations, and *N*_*T*_ and *N*_*C*_ the sample sizes of the music-exposed (treatment) and control groups, respectively.

##### Transformation of proportional outcomes

Some behavioral outcomes were proportions bounded between 0 and 1 (e.g., sucrose preference, proportion of time immobile). Because the variance of proportional data decreases as the mean approaches the bounds, we stabilized variances prior to effect size calculation using the arcsine square-root transformation and the delta method (Macartney et al., 2022). Specifically, for a proportion *M* with standard deviation *SD*, we calculated

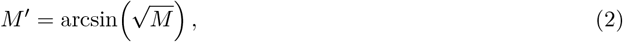

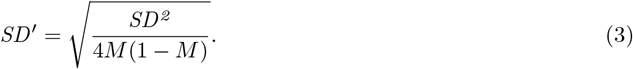

The transformed mean (*M* ^*′*^) and standard deviation (*SD*^*′*^) were then substituted for *M* and *SD* in subsequent calculations of ln *RR* and its sampling variance.

##### Sampling variance and dependence structure

The sampling variance of ln *RR* was calculated as

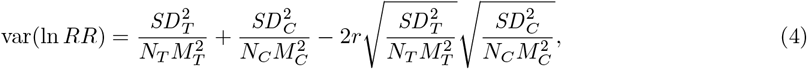

where *SD*_*T*_ and *SD*_*C*_ are the sample standard deviations, *M*_*T*_ and *M*_*C*_ denote the sample means, and *N*_*T*_ and *N*_*C*_ are the sample sizes, of the treatment and control groups, respectively. The parameter *r* represents the correlation between treatment and control responses within the same experimental units. For independent-group designs, *r* was set to zero. For paired or repeated-measures designs, where the same animals contributed to both conditions, *r* was assumed to be 0.5, a conservative value commonly adopted when correlations are not reported (Noble et al., 2017).

#### 2.5.2 Log Variability Ratio (ln *V R*)

Differences in behavioral variability were quantified using the (bias-corrected) log variability ratio (Nakagawa et al., 2015; Senior et al., 2020). Proportional outcomes were excluded because variance is mathematically constrained by the mean in bounded data, and mean shifts can mechanically induce variance shifts (mean– variance dependence).

For continuous outcomes, the log variability ratio was calculated using the bias-corrected estimator

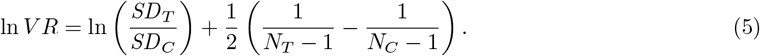

where *N*_*T*_ and *N*_*C*_ are the sample sizes, and *SD*_*T*_ and *SD*_*C*_ are the standard deviations of the treatment (music-exposed) and control groups, respectively.

The corresponding sampling variance was approximated as

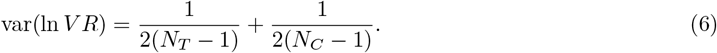

where *N*_*T*_ and *N*_*C*_ denote the treatment and control sample sizes. A positive ln *V R* indicates greater behavioral variability in the music-exposed group, whereas a negative ln *V R* indicates reduced variability relative to controls. For paired or repeated-measures designs, we assumed a within-individual correlation of *r* = 0.5 and used the first-order Taylor expansion (Senior et al., 2020)

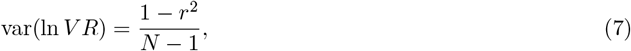

where *N* denotes the number of paired observations contributing to both treatment and control conditions.

We prioritized ln *V R* over the log coefficient of variation ratio (ln *CV R*) to assess absolute rather than relative variability. Although ln *CV R* can partially adjust for mean–variance scaling, it assumes approximately linear scaling of standard deviation with the mean and can become unstable for outcomes with low baseline values. For completeness, analyzes using ln *CV R* are reported in the online Supplementary Material (Ortega et al., 2026).

### 2.6 Meta-analytic models

We fitted multilevel meta-analytic models separately for ln *RR* and ln *V R* using restricted maximum likelihood (REML) estimation in `metafor` (Viechtbauer, 2010). Because individual studies often contributed multiple, non-independent effect sizes (e.g., multiple outcomes, assays, or contrasts from the same experimental cohort), we explicitly modeled dependence among effect sizes in two ways. First, sampling covariances were incorporated using a variance–covariance matrix (**V**) constructed at the cohort level, such that effect sizes derived from the same cohort were assumed to have correlated sampling errors (with *ρ* = 0.5 when within-cohort correlations were not reported), yielding a block-diagonal-structured **V** matrix (Nakagawa et al., 2022; Noble et al., 2017). Second, we included random intercepts for study ID to account for between-study heterogeneity, for effect size ID to accommodate residual within-study variation among effect sizes beyond sampling error, and for strain to control for systematic differences among rodent strains. Moderator analyzes were conducted as separate uni-moderator meta-regressions using the same random-effects structure and the same **V** matrix as the corresponding overall model. Inference was based on t-tests as implemented in `metafor`, and all models were fitted using REML.

### 2.7 Quantifying heterogeneity

Heterogeneity in effect sizes was quantified using the *I*^2^ statistic, which represents the proportion of total observed variance attributable to true heterogeneity rather than sampling error (Lajeunesse, 2011). For multilevel meta-analytic models, a convenient definition of total heterogeneity—equivalent to the standard *I*^2^ used in conventional meta-analyzes (**??**) is given by

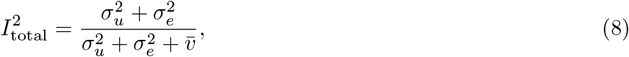

where 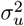 denotes the between-study variance component, 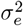 denotes the within-study (effect-size-level) variance component, and 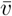 is a representative sampling variance (derived from the sampling variances of individual effect sizes) (Nakagawa and Santos, 2012)

This decomposition further allows heterogeneity to be partitioned into between-study and within-study components:

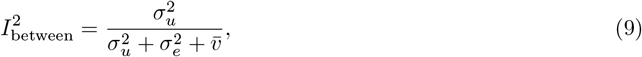

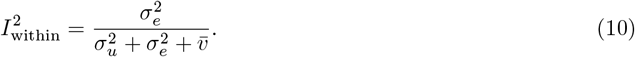

These component-specific *I*^2^ values indicate whether heterogeneity is driven primarily by differences among studies or by residual variation among effect sizes within studies (e.g., due to multiple outcomes, assays, or contrasts derived from the same experiment). All *I*^2^ statistics were treated as descriptive diagnostics rather than inferential quantities.

### 2.8 Small-study effects, time-lag patterns, and sensitivity analyzes

We assessed potential small-study effects and time-lag patterns using complementary visual and model-based approaches appropriate for multilevel meta-analyzes (Nakagawa et al., 2022). These analyzes were treated as diagnostic tools and interpreted cautiously given the substantial heterogeneity and non-independence of effect sizes. First, we visually inspected funnel plots for asymmetry using effect sizes from the fitted multilevel meta-analytic model, plotted against measures of precision (standard error and inverse standard error). Visual inspection was used to assess whether less precise studies showed systematic deviations from the overall pattern. Second, we evaluated small-study effects using an Egger-type regression adapted for multilevel meta-analytic models. In classical Egger’s regression, effect sizes are regressed on their sampling standard errors (Egger et al., 1997). However, for the log response ratio (ln *RR*), effect sizes and sampling variances are inherently correlated, which can bias inference. To avoid this issue, we used the effective sample size:

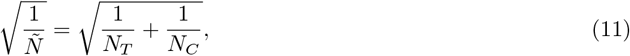

where *N*_*T*_ and *N*_*C*_ are the treatment and control sample sizes, respectively. This term was included as a moderator in multilevel meta-regression models with the same random-effects structure as the primary analyzes. Third, we examined potential time-lag (decline) effects by including mean-centered publication year as a moderator. To disentangle temporal trends from small-study effects, we fitted both uni-moderator models (including either publication year or 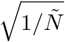) and a multi-moderator model including both predictors simultaneously. All models followed the same variance–covariance structure and random-effects specification as the primary meta-analyzes (Nakagawa et al., 2022). Finally, we assessed the influence of individual studies using a leave-one-out sensitivity analysis. Each study was sequentially removed from the dataset, and the multilevel meta-analytic model was refitted to evaluate the stability of the overall effect estimate. This analysis was used to identify whether any single study exerted disproportionate influence on the estimated mean effect or its uncertainty (Nakagawa et al., 2023b).

### 2.9 Statistical computing environment and software

All analyzes and visualizations were conducted in `R` (R Core Team, 2025) (version 4.5.2) using `RStudio` (Posit team, 2026) (version 2026.01.0+392). We fitted the meta-analytic models using `metafor` (Viechtbauer, 2010)(version 4.8-0). Orchard plots were generated using `orchaRd` (Nakagawa et al., 2023a) (version 2.1.3), figures were constructed with `ggplot2` (Wickham, 2011) (version 4.0.1) and combined using `patchwork` (Pendersen, 2025) (version 1.3.2). Data preprocessing used `tidyverse` (Wickham et al., 2019) (version 2.0.0). *Post hoc* comparisons among moderator levels were performed using `multcomp` (Bretz et al., 2010) (version 1.4-29). All data processing, effect size calculations (ln *RR* and ln *V R*), model specifications, diagnostic checks, bias assessments, and full moderator outputs are provided in a fully reproducible online supplementary document available at: https://santiago-0rtega.github.io/Music_on_anxiety_depression_meta_analysis/. The website includes annotated R code, complete model summaries, and figures.

## 3 Results

### 3.1 Data overview

Twenty studies met the predefined inclusion criteria (Milbratz de Camargo et al., 2017; Camargo et al., 2013; Chen et al., 2019; Cheng et al., 2024; Chikahisa et al., 2007; Niehues da Cruz et al., 2011; Escribano et al., 2014; Flores-Gutiérrez et al., 2018; Freitas Oliveira et al., 2020; Fu et al., 2023, 2025; Krishnamurthy and Rao, 2025; Li et al., 2010; Pangemanan et al., 2024; Papadakakis et al., 2019; Ren and Lu, 2024; Rizzolo et al., 2021; Saghari et al., 2021; Sampaio et al., 2017; Uşak et al., 2020), yielding 298 effect sizes for metaanalysis. Study selection and exclusion decisions are summarized in the PRISMA flow diagram (Fig. S1). Group sample sizes were generally modest, averaging *n* = 9.76 in control groups (SD = 2.53) and *n* = 9.81 in music-exposed groups (SD = 2.59), with mean group sizes across studies ranging from 6.0 to 14.5 animals per group. Most effect sizes were derived from anxiety-like outcomes (77%, *n* = 231), with the remainder from depression-like assays. Effect sizes were nearly evenly split between mice (*Mus musculus*, 48%, four strains) and rats (*Rattus norvegicus*, 52%, two strains; Fig. 1A). Tested animals were predominantly male (60%), followed by female (38%) and mixed-sex cohorts (2%). Note that depression outcomes were almost exclusively evaluated in males, i.e., almost no data for females. The age distribution was strongly skewed toward young adults (61%), with relatively few effect sizes derived from juvenile or older animals. Music exposure was most often acute (≤ 7 days; 50%), and the Elevated Plus Maze was the most frequently used behavioral assay, followed by the Open Field Test (Fig. 1B). Only approximately 18% of effect sizes were derived from animals in which anxiety- or depression-like states were induced using standardized procedures (e.g., chronic unpredictable mild stress).

**Figure 1:**
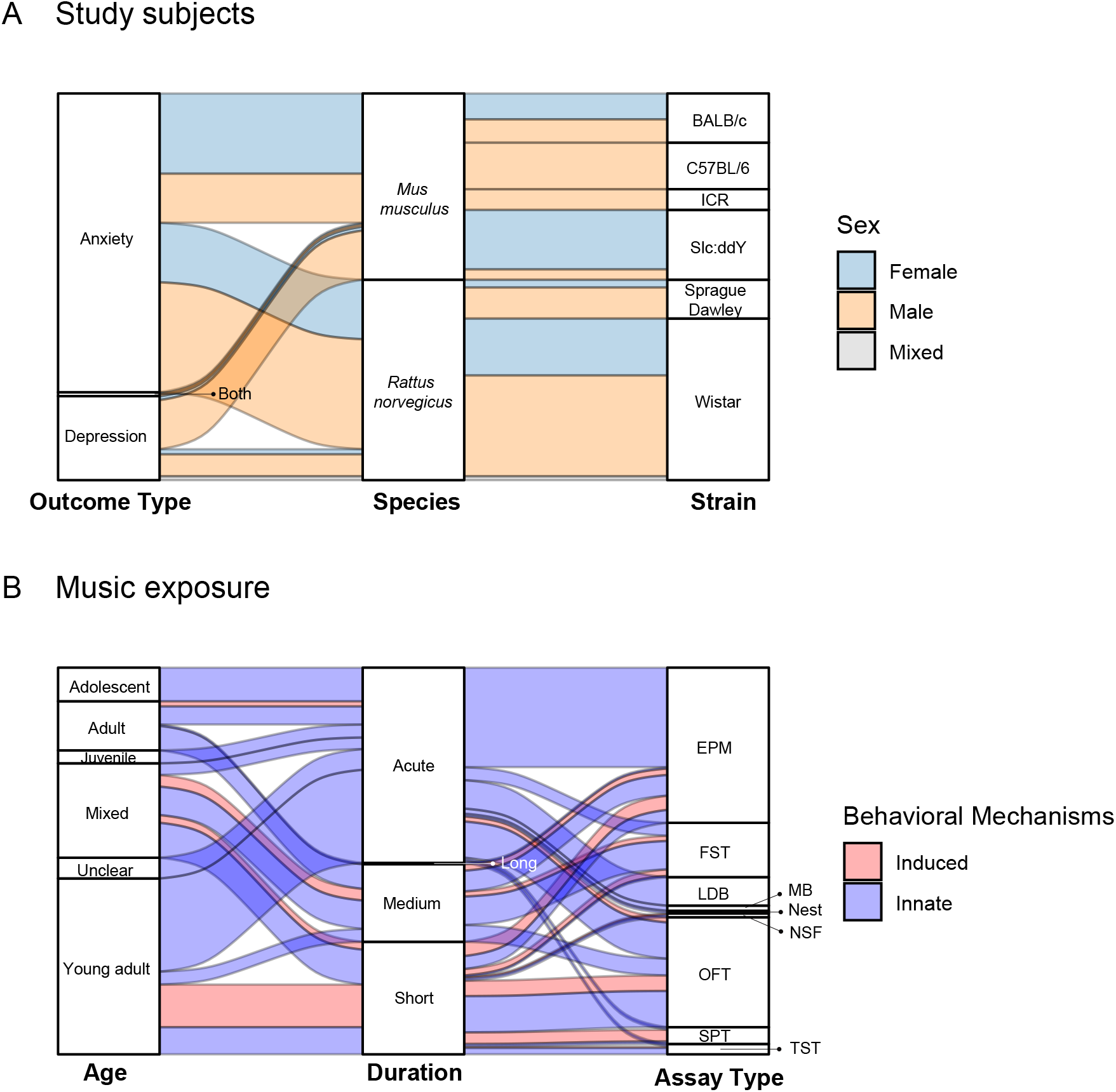
Alluvial plots summarizing the distribution of study characteristics and experimental designs across the included literature. (**A**) Study subjects, showing the flow of effect sizes across outcome type (anxiety-like, depression-like, or both), species (*Mus musculus, Rattus norvegicus*), and strain, with colors indicating sex (female, male, or mixed). (**B**) Music exposure and behavioral assessment, showing the flow of effect sizes across age at exposure, exposure duration, and behavioral assay, with colors indicating behavioral mechanism (innate vs. induced). In panel B, innate assays refer to tests that quantify spontaneous, unconditioned anxiety- or depression-like responses, whereas induced assays refer to tests in which the behavioral response is elicited after an experimental challenge or manipulation (e.g., stress induction). Behavioral assays are abbreviated as follows: EPM = Elevated Plus Maze; OFT = Open Field Test; LDB = Light–Dark Box; NSF = Novelty-Suppressed Feeding; FST = Forced Swim Test; TST = Tail Suspension Test; SPT = Sucrose Preference Test; MB = Marble Burying; Nest = Nest-building test. Flows are proportional to the number of effect sizes contributing to each combination of categories (n = 298).

### 3.2 Overall effects of music exposure

Across all outcomes, music exposure was associated with a significant improvement in mean behavioral outcomes (Fig. 2A). The pooled *lnRR* indicated a 18.7% reduction in anxiety- and depression-like behaviors relative to controls (*β*_Overall *lnRR*_ = −0.208, 95% CI [−0.336, −0.079], *t*_[df = 297]_ = −3.188, *p* = 0.001). Heterogeneity was high 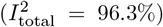, driven primarily by within-study variation among effect sizes 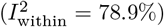, with smaller contributions from between-study heterogeneity 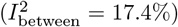. In contrast, variance attributable to strain was negligible relative to other sources (*<* 0.01% of total variance). In contrast, music exposure did not produce a consistent change in behavioral variability (Fig. 2B). Based on analyzes excluding proportional outcomes (*K* = 222), the overall *lnV R* was positive but not significant (*β*_Overall *lnV R*_ = 0.134, 95%*CI*, [−0.118, 0.387], *t*_[df = 221]_ = 1.049, *p* = 0.295). As with *lnRR*, heterogeneity in *lnV R* was high 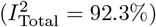 and dominated by within-study variation 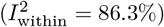, with limited between-study heterogeneity 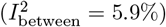. Variance attributable to rodent strain identity was negligible relative to other sources (*<* 0.01% of total variance).

**Figure 2:**
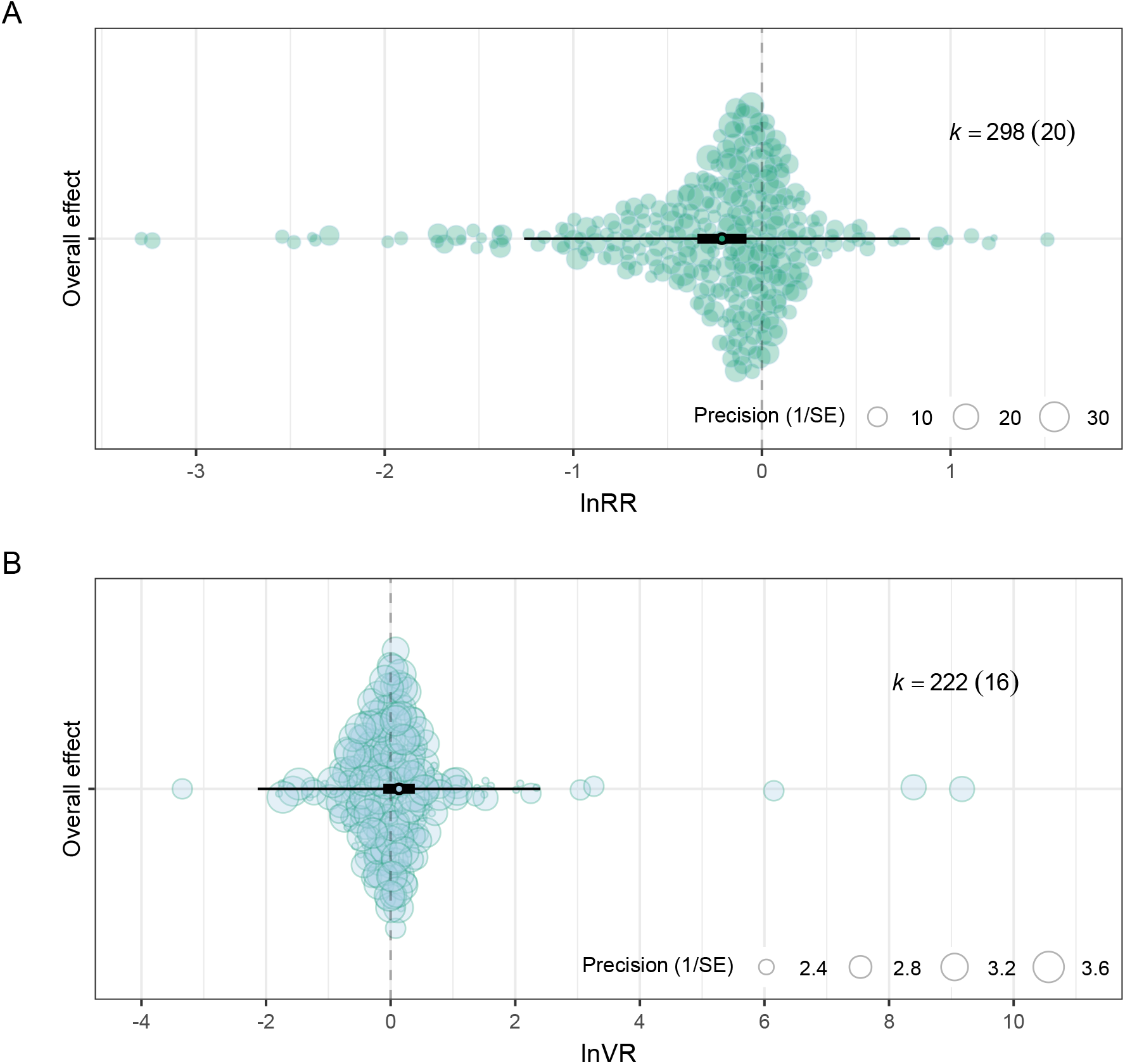
Overall effects of music exposure on behavioral outcomes in rodents. Shown are the pooled estimates of the log response ratio (ln *RR*), quantifying differences in mean anxiety- and depression-like behavior, and the log variability ratio (ln *V R*), quantifying differences in behavioral variability between music-exposed and control groups. Each circle represents an individual effect size, with circle size proportional to precision (inverse of the standard error). The central marker indicates the pooled meta-analytic estimate. Thick and thin horizontal lines represent the 95% confidence interval and 95% prediction interval around the pooled estimate, respectively. Values of *k* indicate the number of effect sizes included in each model (with the value in parentheses indicating the number of studies). Sample sizes differ between panels because proportional outcomes (e.g., bounded measures) were retained in the ln *RR* analyses of mean differences but excluded from the ln *V R* analyses, where variance-based effect sizes are not directly comparable or appropriate for proportional data.

### 3.3 Outcome type: anxiety-like vs. depression-like behaviors

Outcome type did not significantly moderate mean effects of music exposure (Fig. 3A). Music exposure reduced anxiety- and depression-like behaviors on average, and the estimated effect did not differ between outcome types, as indicated by a non-significant anxiety–depression contrast (*β*_Depression-Anxiety_ = 0.002, 95% CI [−0.173, 0.178], *t*_[df=293]_ = 0.025, *p* = 0.979). Conversely, outcome type strongly influenced variability of responses. Anxiety-like outcomes showed increased variability under music exposure (*β*_Anxiety_ = 0.311, 95% CI [0.012, 0.611], *t*_[df=217]_ = 2.047, *p* = 0.042), whereas depression-like outcomes exhibited significantly lower variability relative to anxiety-like assays (*β*_Depression–Anxiety_ = −0.640, 95% CI [−1.078, −0.202], *t*_[df=217]_ = −2.882, *p* = 0.004, Fig. 3B).

**Figure 3:**
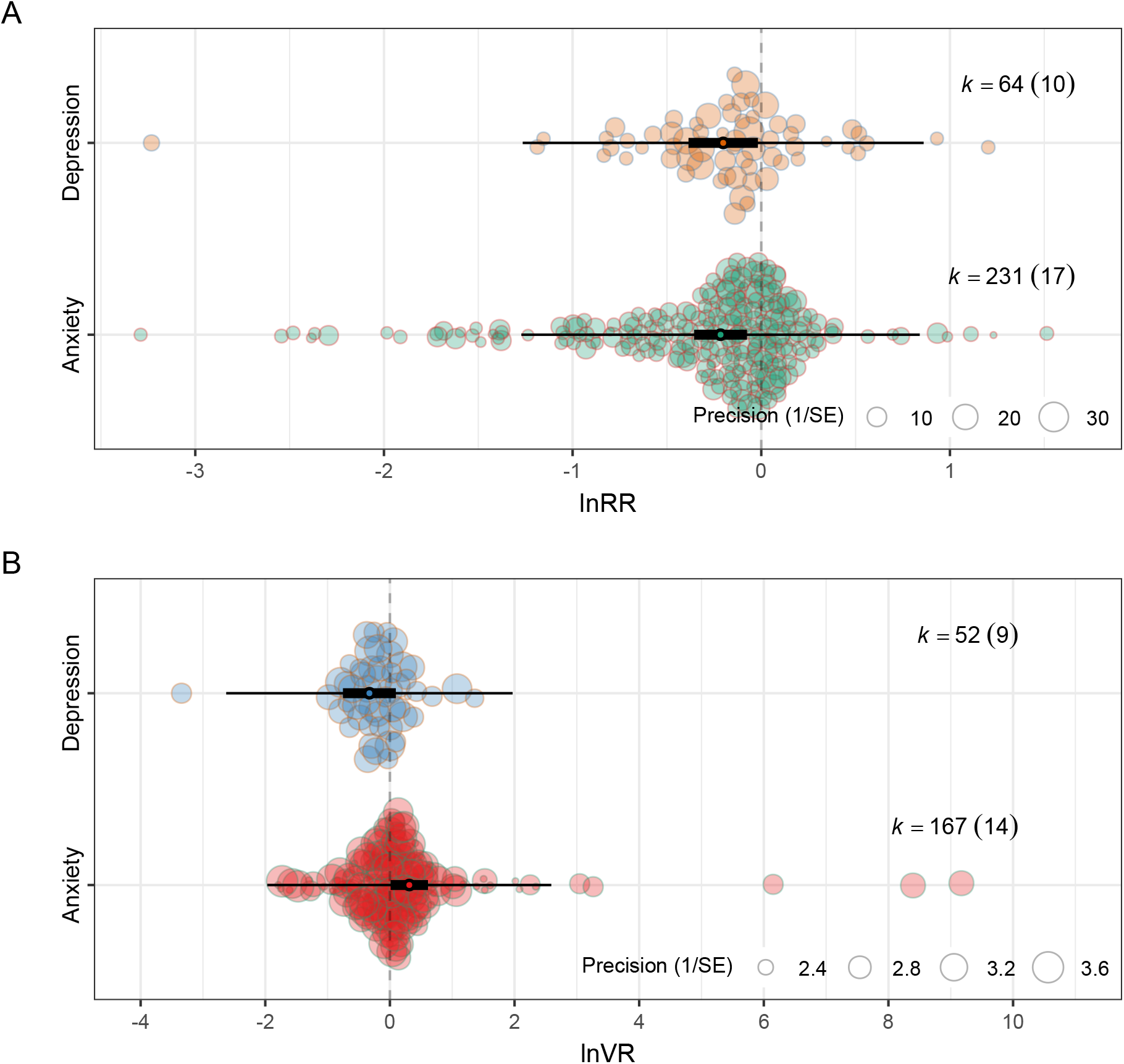
Outcome-type–specific effects of music exposure on rodent behavior based on uni-moderator meta-analyses. (**A**) Effects on mean anxiety-like and depression-like behavioral outcomes quantified using the log response ratio (ln *RR*). (**B**) Effects on behavioral variability in responses quantified using the log variability ratio (ln *V R*). Each circle represents an individual effect size, with circle size proportional to precision (inverse of the standard error). Thick and thin horizontal lines represent 95% confidence and prediction intervals around the pooled estimate, respectively. Effect sizes classified as testing both outcome types were excluded due to low sample size (*k <* 5), and proportional outcomes were excluded from the ln *V R* analyses.

### 3.4 Uni-moderator analyzes of mean behavioral effects (ln *RR*)

Uni-variate moderator analyzes were conducted on effect sizes pooled across outcome types to identify sources of heterogeneity in mean behavioral responses. Most moderators explained modest proportions of heterogeneity, with marginal *R*^2^ values generally below 5% (Table 2). Behavioral assay accounted for the largest share of heterogeneity (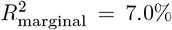, Fig. 4A). The Light–Dark Box yielded more negative ln *RR* values (i.e., stronger reductions in anxiety- or depression-like behavior) than both the Elevated Plus Maze (*β*_LDB–EPM_ = −0.272, *z* = −2.037, *p* = 0.04) and the Open Field Test (*β*_OFT-LDB_ = 0.485, *z* = 3.232, *p* = 0.001). The Open Field Test, in turn, showed more positive ln *RR* values than the Forced Swim Test (*β*_OFT–FST_ = 0.213, *z* = 2.618, *p* = 0.008) and the Elevated Plus Maze (*β*_OFT–EPM_ = 0.246, *z* = 2.112, *p* = 0.034). Life stage at exposure explained 4.1% of heterogeneity (Fig. 4B). Mean effects were negative across all life stages. Control condition accounted for 3.3% of heterogeneity (Fig. 4C), with white-noise controls associated with larger reductions in anxiety- and depression-like behaviors than ambient sound controls (*β*_White noise–Ambient sound_ = −0.261, 95% CI [−0.458, −0.064], *t*_296_ = −2.616, *p* = 0.009). Behavioral mechanism explained 5.4% of heterogeneity (Fig. 4D); effect sizes derived from innate behavioral assays showed more positive ln *RR* values (i.e., smaller reductions) than those from induced protocols (*β*_Innate–Induced_ = 0.320, 95% CI [0.106, 0.533], *t*_296_ = 2.949, *p* = 0.003). All remaining moderators (sex, exposure duration, music meta-genre, experimental design, and relative timing) each explained ≤ 2.3% of heterogeneity. Nevertheless, relative timing of exposure 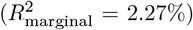 showed a significant contrast: music presented concurrently with behavioral testing was associated with more positive ln *RR* values (i.e., weaker reductions) than exposure occurring before testing (*β*_Concurrent–Before_ = 0.557, *z* = 2.299, *p* = 0.021) or both before and after testing (*β*_Concurrent–Both_ = 0.529, *z* = 2.125, *p* = 0.033).

**Table 2:** Proportion of total heterogeneity (marginal *R*^2^) explained by individual moderators. Values represent the proportion of heterogeneity explained by each moderator in separate uni-variate models fitted to effect sizes pooled across anxiety- and depression-like outcomes. These estimates should not be interpreted as independent or additive contributions.

| **Moderator** | **lnRR Marginal $R^2$** | **lnVR Marginal $R^2$** |
| --- | --- | --- |
| Behavioral assay | 7.17% | 11.19% |
| Behavioral mechanism | 5.42% | 0.22% |
| Age at exposure | 4.16% | 3.88% |
| Control condition | 3.30% | 0.23% |
| Relative timing | 2.27% | 0.55% |
| Music meta-genre | 2.19% | 8.41% |
| Experimental design | 1.99% | 5.6% |
| Sex | 2.07% | 0.30% |
| Exposure duration | 2.16% | 1.01% |
| Experimental procedure | 0.50% | 0.22% |
| Outcome type | 0.00% | 5.37% |

**Figure 4:**
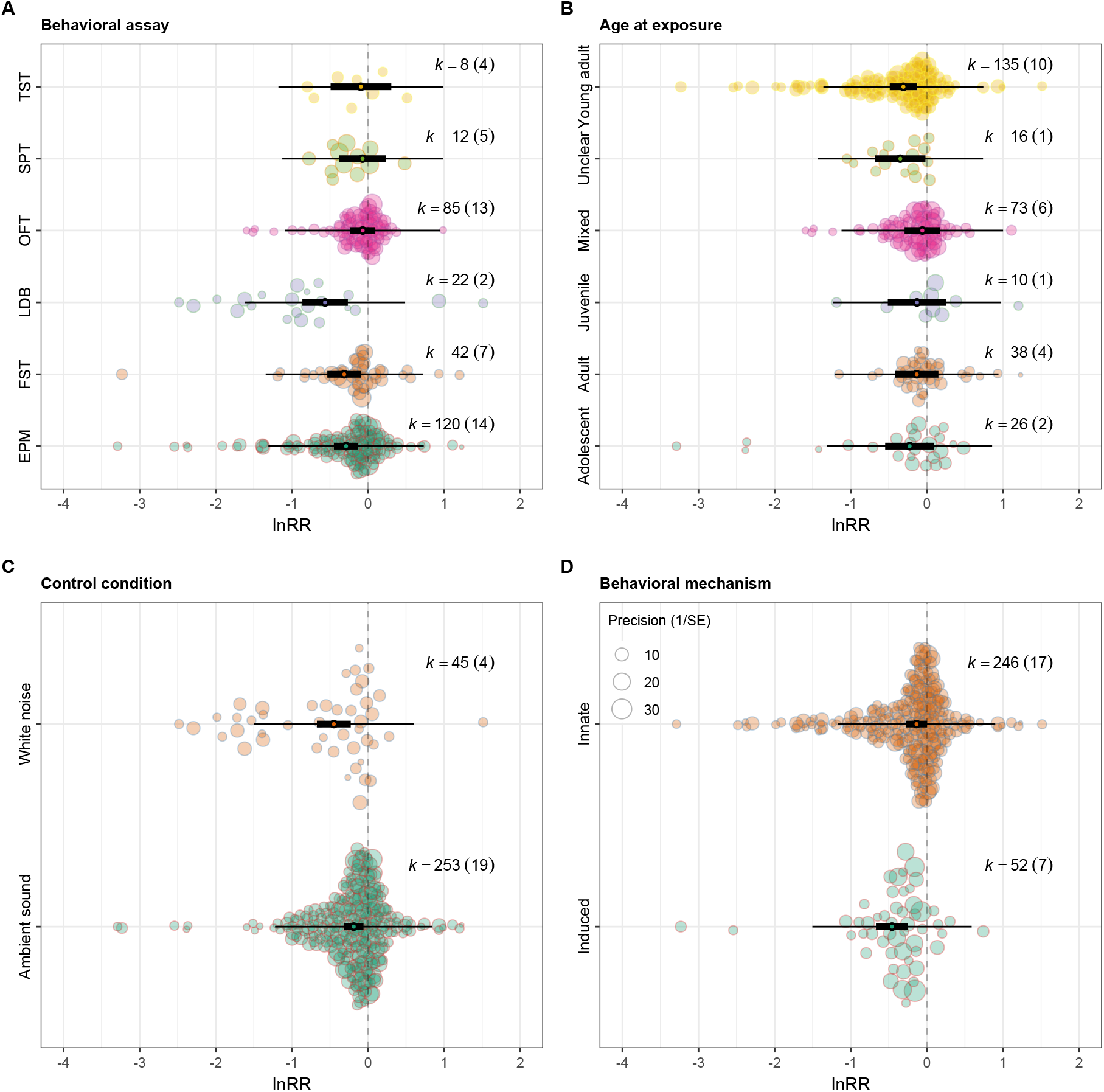
Uni-moderator analyzes of mean behavioral effects of music exposure (ln *RR*). Orchard plots show pooled estimates from multilevel meta-analytic models for selected moderators, with effect sizes pooled across anxiety- and depression-like outcomes. Points represent individual effect sizes, with point size proportional to precision; thick horizontal lines indicate 95% confidence intervals, and thin horizontal lines indicate 95% prediction intervals. Negative values indicate beneficial effects of music exposure (i.e., reduced anxiety- or depression-like behavior). Panels show moderators for (A) behavioral assay, (B) life stage at exposure, (C) control condition type, and (D) behavioral mechanism (induced vs. innate). Behavioral assays are abbreviated as follows: Elevated Plus Maze (EPM), Open Field Test (OFT), Light–Dark Box (LDB), Forced Swim Test (FST), Tail Suspension Test (TST), and Sucrose Preference Test (SPT).

### 3.5 Uni-moderator analyzes of behavioral variability (ln *V R*)

Uni-moderator analyses of ln *V R* examined how music exposure influenced behavioral consistency relative to controls. Negative ln *V R* values indicate greater consistency (reduced dispersion across individuals), whereas positive values indicate increased dispersion and less predictable responses. Overall, most moderators explained relatively small proportions of heterogeneity in ln *V R* (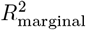 typically ≤ 6%). Behavioral assay was the strongest moderator of variability (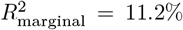 Fig. 5A). Relative to the Elevated Plus Maze (EPM), music exposure was associated with significantly lower variability in the Forced Swim Test (*β*_FST–EPM_ = −0.906, *z* = −3.375, *p <* 0.001), Open Field Test (*β*_OFT–EPM_ = −0.755, *z* = −3.369, *p <* 0.001), and Tail Suspension Test (*β*_TST–EPM_ = −1.251, *z* = −2.713, *p* = 0.006), indicating more predictable responses in these assays. The Light–Dark Box showed a marginal reduction in variability relative to the EPM (*β*_LDB–EPM_ = −0.552, *z* = −1.900, *p* = 0.057). Life stage at exposure explained 3.8% of heterogeneity (Fig. 5B). Outcomes from rodents tested across mixed age groups were significantly less variable than those tested during adolescence (*β*_Mixed–Adolescent_ = −0.851, *z* = −2.116, *p* = 0.034). Music meta-genre also accounted for substantial heterogeneity (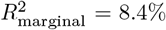 Fig. 5C). Relative to mixed music, exposure to popular contemporary music (*β*_Contemporary–Mixed_ = 1.020, *z* = 2.401, *p* = 0.016) and traditional/folk/world music (*β*_World–Mixed_ = 1.190, *z* = 2.190, *p* = 0.028) was associated with increased variability. In contrast, Western art/orchestral music was associated with significantly lower variability than both contemporary (*β*_Orchestral–Contemporary_ = −0.977, *z* = −2.879, *p* = 0.004) and traditional/folk/-world music (*β*_Orchestral–World_ = −1.147, *z* = −2.404, *p* = 0.016). Experimental design explained 5.6% of heterogeneity (Fig. 5D). Posttest-only designs were associated with greater variability than factorial designs (*β*_Posttest–Factorial_ = 0.623, *z* = 2.869, *p* = 0.004), whereas repeated-measures designs showed lower variability than posttest-only designs (*β*_Repeated–Posttest_ = −0.849, *z* = −2.311, *p* = 0.020). All remaining moderators (sex, exposure duration, control condition, experimental procedures, behavioral mechanism, and relative timing) explained negligible heterogeneity (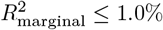) and showed no significant moderating effects.

**Figure 5:**
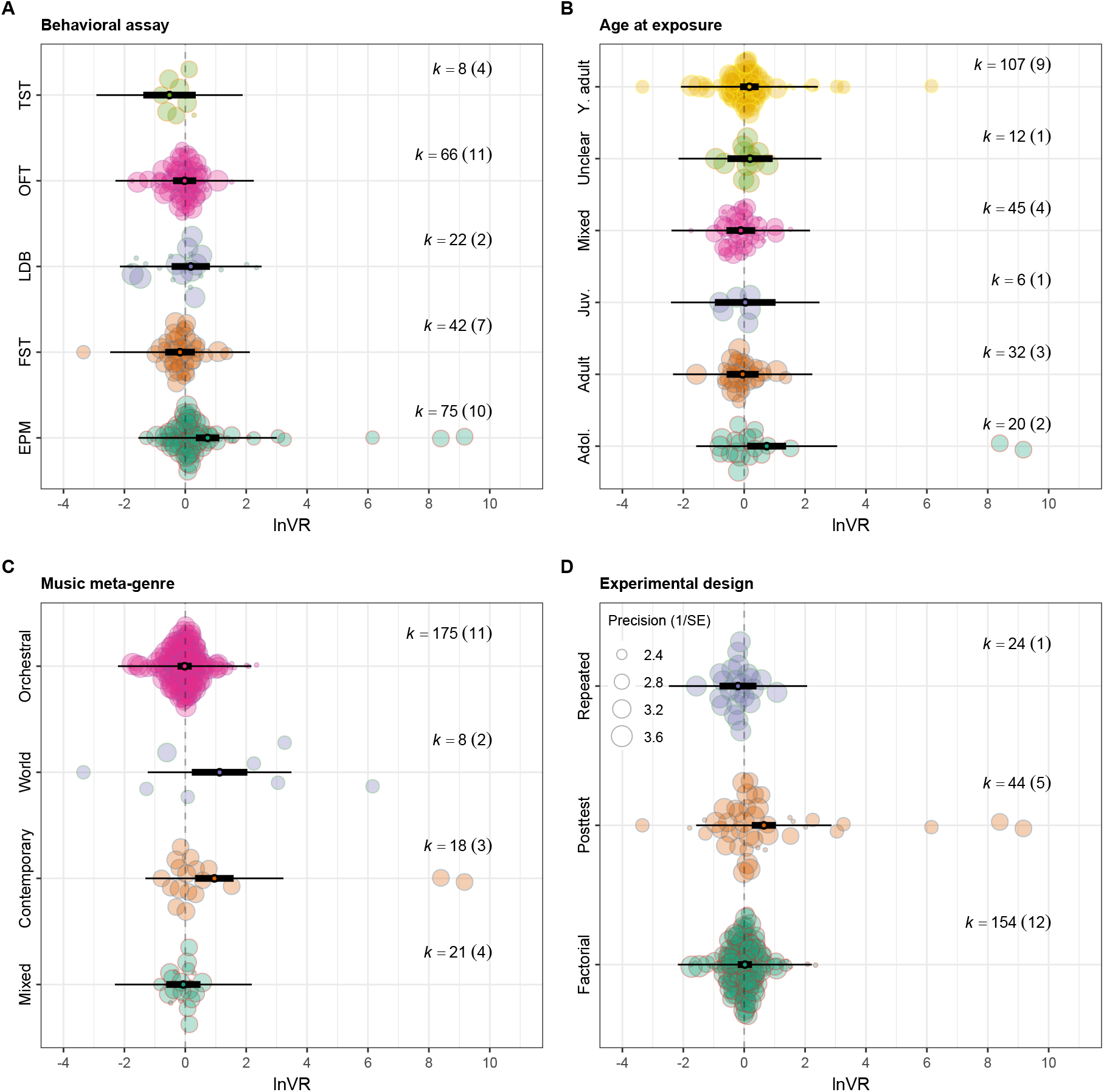
Uni-moderator analyzes of behavioral variability under music exposure (ln *V R*). Orchard plots show pooled estimates from multilevel meta-analytic models for selected moderators, with effect sizes pooled across anxiety- and depression-like outcomes. Points represent individual effect sizes, with point size proportional to precision; thick horizontal lines indicate 95% confidence intervals, and thin horizontal lines indicate 95% prediction intervals. Positive values indicate increased behavioral variability (i.e., less predictable responses), whereas negative values indicate reduced variability (i.e., more predictable responses) following music exposure. Panels show moderators for (A) behavioral assay, (B) age at exposure, (C) music meta-genre, and (D) experimental design. Behavioral assays are abbreviated as follows: Elevated Plus Maze (EPM), Open Field Test (OFT), Light–Dark Box (LDB), Forced Swim Test (FST), Tail Suspension Test (TST), and Sucrose Preference Test (SPT).

### 3.6 Small-study effects, time-lag patterns, and sensitivity analyzes

We found no evidence of small-study effects or time-lag bias. Funnel plots based on model residuals showed no clear asymmetry (Fig. S2), and the Egger’s-type multilevel regression using inverse effective sample size was non-significant (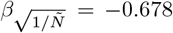, 95% CI [−2.396, 1.03], *t*_296_ = −0.777, *p* = 0.437; Fig. 6A). Publication year did not predict effect size magnitude, either alone or when fitted jointly with the precision term (*β*_*Y ear*_ = −0.004, 95% CI [−0.028, 0.019], *t*_295_ = −0.692, *p* = 0.715; Fig. 6B). Leave-one-out analyzes confirmed the robustness of the overall ln *RR* estimate. Sequential exclusion of individual studies yielded highly consistent pooled estimates, with all confidence intervals overlapping the full-sample estimate and no single study exerting disproportionate influence (Fig. S3).

**Figure 6:**
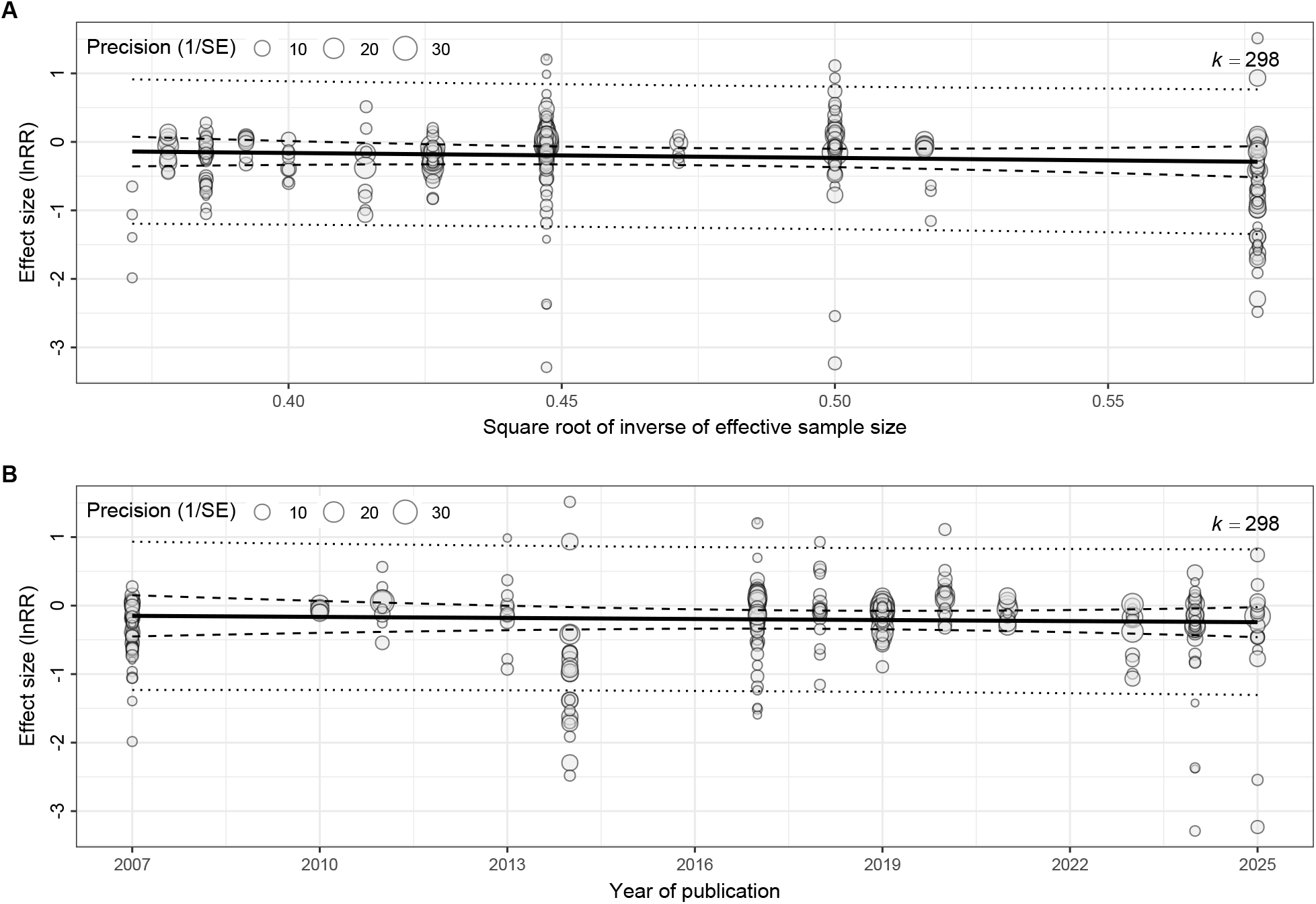
Associations between effect sizes and (A) the square root of the inverse effective sample size and (B) publication year. Point size reflects precision; solid lines show fitted meta-regressions, and dashed and dotted lines indicate 95% confidence and prediction intervals, respectively.

### 3.7 Risk of bias and reporting transparency

Reporting of methodological safeguards and stimulus characteristics was variable across studies. Randomization was reported in 65% of studies, whereas blinding of outcome assessment was reported in only 35%. Reporting of stimulus characteristics (frequency and loudness) was incomplete: music-stimulus details were provided in 55% of studies, but control-conditions characteristics (ambient sound or white noise) were rarely reported (10%). Timing-related information (i.e., whether behavioral testing occurred during the light or dark phase) was reported in 25% of studies. No study explicitly reported using a formal risk-of-bias guideline (0%). Attrition handling was rarely described (10%), and selective outcome reporting was flagged in 20% of studies. These reporting gaps informed the interpretation of heterogeneity and are considered further in the Discussion.

## 4 Discussion

Across experimental contexts, music exposure reduced anxiety- and depression-like behaviors by approximately 18% relative to control conditions, as indicated by a significantly negative overall ln *RR* (Fig. 2A). This reduction occurred across both anxiety- and depression-like outcomes (Fig. 3A) and across life stages at exposure (Fig. 4A), demonstrating that music exposure shifts affect-related behavior in a consistent direction across diverse experimental settings. Although effect sizes showed substantial heterogeneity (Table 2), the predominance of negative effect sizes supports a robust directional effect.

Music exposure did not systematically alter behavioral variability. The overall ln *V R* estimate was close to zero and non-significant (Fig. 2B), indicating that music exposure neither consistently reduced nor increased dispersion across individuals. In other words, music exposure shifted mean affect-related behavior without broadly homogenizing responses or amplifying inter-individual differences. This distinction highlights the conceptual difference between mean effects (ln *RR*), which capture shifts in average behavior, and variability effects (ln *V R*), which capture changes in dispersion (Nakagawa et al., 2015). The absence of a global ln *V R* effect indicates that music alters central tendency without uniformly reshaping between-individual variation. However, moderator analyses revealed structured heterogeneity in variability responses, particularly across behavioral assays, music meta-genre, and experimental design (Fig. 5B–C). Strain contributed negligibly to heterogeneity in both mean and variability models. Although strain differences often influence behavioral outcomes (Russo et al., 2012), the available evidence suggests broadly similar directional effects across the mouse and rat strains represented in this synthesis.

### 4.1 Music exposure in the context of environmental enrichment

The direction of the mean effects aligns with findings from other non-pharmacological interventions, particularly voluntary exercise and environmental enrichment. Exercise reliably reduces immobility in stress-coping assays and increases exploratory behavior in anxiety-related tests (Brenes et al., 2020; Duman et al., 2008; Mul and Mul, 2018; Yang et al., 2025). Although we cannot directly compare effect magnitudes across paradigms, the shared direction of effects suggests that music functions as a sensory form of environmental enrichment rather than as a pharmacological analogue. Music exposure produced stronger reductions in induced paradigms than in assays measuring baseline affective behavior (Fig. 4D). Induced models (e.g., chronic stress or learned helplessness paradigms) elevate anxiety- and depression-like phenotypes above typical levels (Mineur et al., 2006), thereby creating greater scope for behavioral improvement. Interventions that modulate stress physiology would therefore be expected to produce larger effects under heightened disturbance than under baseline conditions, where responses may already fall within a more restricted range.

Timing also influenced effect magnitude. Studies that delivered music prior to behavioral testing showed larger reductions in anxiety- and depression-like behaviors than studies presenting music concurrently with testing. Rather than contradicting evidence for acute music-related changes in arousal or cognition (Rickard et al., 2005), this pattern suggests that music may primarily shift baseline affective state rather than enhance task performance. When introduced during testing, music may act as an additional sensory stimulus that interacts with task demands, whereas prior exposure allows physiological and affective modulation to occur before assessment. Several mechanisms may account for these patterns. Music may modulate stress physiology through the hypothalamic–pituitary–adrenal (HPA) axis, reducing stress reactivity and downstream glucocorticoid signaling (Smith and Vale, 2006; **?**). Alternatively, music may increase environmental predictability. Chronic unpredictable mild stress reliably induces anxiety- and depression-like phenotypes (Willner, 2017), whereas music exposure typically occurs in stable, repeatable contexts. By increasing sensory predictability, music may dampen stress responses associated with uncertainty. This interpretation aligns with the joint ln *RR*–ln *V R* pattern: music consistently shifts mean behavior while altering variability only in specific contexts. As with other enrichment-based interventions, heterogeneity is therefore expected rather than anomalous (Simpson and Kelly, 2011; Würbel, 2001).

### 4.2 Assay-, stimulus-, and design-specific variability

Behavioral assay strongly moderated variability responses. Relative to the Elevated Plus Maze (EPM), music exposure reduced variability in the Forced Swim Test and Tail Suspension Test (Fig. 5A). These assays constrain animals to a limited behavioral repertoire and probe stress-coping states (Cryan and Holmes, 2005; Leite-Almeida et al., 2022). Under such constraints, music appears to promote convergence toward similar coping responses, increasing predictability across individuals. In contrast, music exposure increased variability in the EPM. The EPM assesses approach–avoidance conflict and typically produces substantial between-individual variability (Albani et al., 2015; Knight et al., 2021; Walf and Frye, 2007). Music may amplify pre-existing differences in how animals balance exploration and avoidance, increasing dispersion even when mean anxiety-like behavior declines. Increased variability in this context does not imply adverse effects; rather, it reflects sensitivity of this behavioral domain to individual traits and assay structure. These findings carry practical implications. Researchers who prioritize reproducibility and precise estimation of mean effects may obtain more consistent outcomes using assays that exhibited lower variability under music exposure, such as the Forced Swim Test or Tail Suspension Test. In contrast, assays like the EPM may require larger samples or explicit modeling of heterogeneity to achieve comparable precision. No assay is inherently superior; instead, tasks differ in how consistently animals respond under identical auditory conditions.

Music meta-genre also structured variability. Studies using popular contemporary and traditional or folk music showed greater dispersion than studies using mixed stimuli, whereas Western art or orchestral music did not differ significantly from the reference category (Fig. 5C). Although genre categories are coarse, they likely capture differences in auditory temporal structure and predictability. Predictable auditory patterns tend to elicit more consistent neural responses (Auksztulewicz et al., 2017; Heilbron and Chait, 2018), whereas less predictable rhythms amplify differential processing and expectation violations (Foldal et al., 2020). Thus, genre-dependent variability likely reflects differences in how consistently animals encode and integrate auditory input. Experimental design also influenced variability. Posttest-only designs produced greater dispersion than factorial or repeated-measures designs (Fig. 5D). Structured designs reduce unexplained variance by controlling baseline differences, thereby yielding more stable effect estimates. Consequently, some variability attributed to music exposure likely reflects differences in design architecture rather than biological effects alone.

### 4.3 Heterogeneity, reporting practices, and robustness

The combined ln *RR* and ln *V R* results highlight pronounced heterogeneity in music exposure protocols across studies (Kühlmann et al., 2018; Le et al., 2025). Differences in genre, exposure duration, timing, acoustic properties, and experimental context are likely to interact in non-additive ways, complicating mechanistic inference. Notably, music meta-genre explained little heterogeneity in mean effects but substantial heterogeneity in behavioral variability, indicating that average outcomes alone can obscure important dimensions of behavioral sensitivity. From this perspective, heterogeneity largely reflects diversity in how music exposure is implemented and experienced rather than inconsistency in effect direction.

Interpretation of results is further constrained by variable reporting practices. Although randomization was commonly reported, statements of blinding, attrition handling, and detailed characterization of auditory stimuli and control conditions were often absent, limiting the ability to isolate music-specific effects from broader sensory or procedural influences. Despite these limitations, multiple diagnostics support the robustness of the main conclusions of our meta-analysis: we detected no evidence of small-study effects or time-lag bias, and leave-one-out analyses showed that the overall mean effect was not driven by any single study. Thus, the observed heterogeneity is more consistent with biological and methodological diversity than with systematic bias.

Importantly, the available evidence supports generalization primarily within the range of experimental conditions represented in the literature. Most studies involved young adult laboratory mice or rats used relatively short-term music exposures under controlled housing conditions. Accordingly, the estimated ≈ 19% reduction in anxiety- and depression-like behaviors should be interpreted as applicable to similar laboratory contexts. Extrapolation to other developmental stages, longer-term exposures, different species, or more complex environmental settings should be made cautiously until directly tested.

An additional under-reported source of heterogeneity concerns circadian timing. Rodents are nocturnal, and affect-related behaviors can vary with time of day; although habituation to testing procedures may attenuate strong circadian effects, explicit reporting of whether music exposure and behavioral testing occurred during the light or dark phase was uncommon. This suboptimal reporting limits assessment of whether circadian context contributed to unexplained heterogeneity in either mean effects or behavioral variability and may obscure subtle interactions between auditory stimulation and task sensitivity (Saré et al., 2021).

Most primary studies were also not designed to detect changes in behavioral variability, as sample sizes and analytical frameworks typically prioritize mean differences between groups. Consequently, the context-dependent ln *V R* patterns identified here likely represent conservative estimates of how music exposure influences behavioral dispersion. That structured variability effects emerged despite these constraints suggests that music can differentially shape individual responses under specific behavioral contexts.

A further pervasive design limitation concerns the unit of replication. None of the included studies explicitly treated cage, rather than individual animal, as the experimental unit, nor did they report statistical control for cage effects (e.g., inclusion of cage as a random effect). Because socially housed rodents within the same cage are not statistically independent, failure to account for this structure can result in pseudo-replication and inflated precision (Lazic, 2010). This issue is particularly relevant given the widespread reliance on t-tests or ANOVA-based analyses, with limited use of hierarchical or mixed-effects modeling. Such practices are common in preclinical research but are increasingly recognized as contributors to reduced reproducibility (Aarts et al., 2014; Holman et al., 2015; Voelkl et al., 2020).

### 4.4 Study-specific limitations and recommendations for future research

Several limitations of the present synthesis point to clear priorities for future work on music exposure and affect-related behavior. Greater standardization and reporting of music exposure protocols—including stimulus selection, exposure schedule, and acoustic parameters—would improve comparability across studies. Explicit reporting of temporal context, including light–dark phase and habituation procedures, is also essential for evaluating potential circadian contributions. Finally, future studies should more directly address behavioral variability through larger sample sizes, repeated-measures designs, or analyzes that retain individual-level response distributions rather than relying solely on group means. Future work would also benefit from explicitly modeling hierarchical structure (e.g., cage or cohort effects) at the design and analysis stages.

### 4.5 Conclusions

This meta-analysis shows that music exposure is associated, on average, with an approximately 18% reduction in anxiety- and depression-like behaviors in rodents, while effects on behavioral variability are strongly context dependent. Music exposure primarily shifts the central tendency of affect-related behavior, with changes in predictability emerging only under specific behavioral and stimulus conditions. Joint consideration of mean effects and variability therefore provides a more complete framework for interpreting heterogeneous experimental results and for evaluating sensory and environmental interventions.

## 5 Author contributions

Conceptualization: S.O., S.N., & M.L.; Methodology: M.L., S.N., & S.O.; Validation: A.L, S.N., & M.L.; Formal analysis: S.O.; Investigation: All.; Data curation: S.O.; Writing - Original Draft: S.O.; Writing - Review & Editing : All; Visualization: S.O.; Supervision: S.O., S.N., & M.L.; Project administration: S.N. Funding acquisition: S.N.

## 7 Data availability

All data and code are available at https://github.com/Santiago-0rtega/Music_on_anxiety_depression_meta_analysis

## References

Aarts, E., Verhage, M., Veenvliet, J.V., Dolan, C.V., Van Der Sluis, S., 2014. A solution to dependency: using multilevel analysis to accommodate nested data. Nature neuroscience 17, 491–496. URL: https://pubmed.ncbi.nlm.nih.gov/24671065/, doi:10.1038/nn.3648.

Airaksinen, E., Larsson, M., Lundberg, I., Forsell, Y., 2024. Anxiety and depressive personality disorders in the modern world. Acta Psychologica 246, 104285. URL: https://doi.org/10.1017/s0033291703008559, doi:10.1017/S0033291703008559.

Albani, S.H., Andrawis, M.M., Abella, R.J.H., Fulghum, J.T., Vafamand, N., Dumas, T.C., 2015. Behavior in the elevated plus maze is differentially affected by testing conditions in rats under and over three weeks of age. Frontiers in Behavioral Neuroscience 9, 98316. URL: https://www.frontiersin.org, doi:10.3389/fnbeh.2015.00031.

Arnold, C.A., Bagg, M.K., Harvey, A.R., 2024. The psychophysiology of music-based interventions and the experience of pain. Frontiers in Psychology 15, 1361857. doi:10.3389/fpsyg.2024.1361857.

Auksztulewicz, R., Barascud, N., Cooray, G., Nobre, A.C., Chait, M., Friston, K., 2017. The Cumulative Effects of Predictability on Synaptic Gain in the Auditory Processing Stream. The Journal of neuroscience : the official journal of the Society for Neuroscience 37, 6751–6760. URL: https://pubmed.ncbi.nlm.nih.gov/28607165/, doi:10.1523/JNEUROSCI.0291-17.2017.

Balkwill, L.L., Thompson, W.F., 1999. A cross-cultural investigation of the perception of emotion in music: Psychophysical and cultural cues. Music Perception 17, 43–64. doi:10.2307/40285811.

Boulenger, J.P., 2004. Residual symptoms of depression: clinical and theoretical implications. European Psychiatry 19, 209–213. URL: https://www.cambridge.org/core/journals/european-psychiatry/article/residual-symptoms-of-depression-clinical-and-theoretical-implications/B1A6ADA633B47F661E3E5CBFC510BB98, doi:10.1016/j.eurpsy.2004.04.001.

Brenes, J.C., Fornaguera, J., Sequeira-Cordero, A., 2020. Environmental Enrichment and Physical Exercise Attenuate the Depressive-Like Effects Induced by Social Isolation Stress in Rats. Frontiers in Pharmacology 11, 541083. URL: https://www.frontiersin.org, doi:10.3389/fphar.2020.00804.

Bretz, F., Horthorn, T., Westfall, P., 2010. Multiple Comparisons Using R. Chapman & Hall/CRC. URL: https://www.routledge.com/Multiple-Comparisons-Using-R/Bretz-Hothorn-Westfall/p/book/9781584885740.

Milbratz de Camargo, A., Bonde, H., Delwing Dal Magro, D., Delwing de Lima, D., de Azevedo Campanella, L.C., 2017. Cocoa and classical music: effect on anxiety and antioxidant activity in Wistar rats. Archivos Latinoamericanos de Nutrición 67.

Camargo, A.M.d., Lima, D.D.d., Dal Magro, D.D., Seubert, J.K., Cruz, J.N.d., Cruz, J.G.P.d., 2013. Adjuvant effects of classical music on simvastatin induced reduction of anxiety but not object recognition memory in rats. Psychology & Neuroscience 6, 403–410. URL: https://doi.apa.org/doi/10.3922/j.psns.2013.3.19, doi:10.3922/j.psns.2013.3.19.

Chen, S., Liang, T., Zhou, F.H., Cao, Y., Wang, C., Wang, F.Y., Li, F., Zhou, X.F., Zhang, J.Y., Li, C.Q., 2019. Regular Music Exposure in Juvenile Rats Facilitates Conditioned Fear Extinction and Reduces Anxiety after Foot Shock in Adulthood. BioMed Research International 2019, 1–10. URL: https://www.hindawi.com/journals/bmri/2019/8740674/, doi:10.1155/2019/8740674.

Cheng, H.Y., Xie, H.X., Tang, Q.L., Yi, L.T., Zhu, J.X., 2024. Light and classical music therapies attenuate chronic unpredictable mild stress-induced depression via BDNF signaling pathway in mice. Heliyon 10, e34196. URL: https://linkinghub.elsevier.com/retrieve/pii/S2405844024102277, doi:10.1016/j.heliyon.2024.e34196.

Chikahisa, S., Sano, A., Kitaoka, K., Miyamoto, K.I., Sei, H., 2007. Anxiolytic effect of music depends on ovarian steroid in female mice. Behavioural Brain Research 179, 50–59. URL: https://linkinghub.elsevier.com/retrieve/pii/S0166432807000289, doi:10.1016/j.bbr.2007.01.010.

Niehues da Cruz, J., Delwing de Lima, D., Delwing Dal Magro, D., Geraldo Pereira da Cruz, J., 2011. The Power of Classic Music to Reduce Anxiety in Rats Treated with Simvastatin. Basic and Clinical Neuroscience 2, 5–11. URL: https://bcn.iums.ac.ir/article-1-173-en.html.

Cryan, J.F., Holmes, A., 2005. The ascent of mouse: advances in modelling human depression and anxiety. Nature Reviews Drug Discovery 2005 4:9 4, 775–790. URL: https://www.nature.com/articles/nrd1825, doi:10.1038/nrd1825.

Duman, C.H., Schlesinger, L., Russell, D.S., Duman, R.S., 2008. Voluntary exercise produces antidepressant and anxiolytic behavioral effects in mice. Brain Research 1199, 148–158. URL: https://doi.org/10.1097/00006842-200009000-00006, doi:10.1016/j.brainres.2007.12.047.

Egger, M., Smith, G.D., Schneider, M., Minder, C., 1997. Bias in meta-analysis detected by a simple, graphical test. BMJ 315, 629–634. URL: https://www.bmj.com/content/315/7109/629 https://www.bmj.com/content/315/7109/629.abstract, doi:10.1136/BMJ.315.7109.629.

Escribano, B., Quero, I., Feijóo, M., Tasset, I., Montilla, P., Túnez, I., 2014. Role of noise and music as anxiety modulators: Relationship with ovarian hormones in the rat. Applied Animal Behaviour Science 152, 73–82. URL: https://linkinghub.elsevier.com/retrieve/pii/S0168159113002955, doi:10.1016/j.applanim.2013.12.006.

Fancourt, D., Finn, S., 2019. What is the evidence on the role of the arts in improving health and well-being? Technical Report. WHO Regional Office for Europe. Copenhagen. URL: https://www.ncbi.nlm.nih.gov/books/NBK553773/.

Fancourt, D., Ockelford, A., Belai, A., 2014. The psychoneuroimmunological effects of music: A systematic review and a new model. Brain, Behavior, and Immunity 36, 15–26. URL: https://www.sciencedirect.com/science/article/abs/pii/S0889159113005138?via%3Dihub, doi:10.1016/j.bbi.2013.10.014.

Ferrari, A., 2022. Global, regional, and national burden of 12 mental disorders in 204 countries and territories, 1990–2019: a systematic analysis for the Global Burden of Disease Study 2019. The Lancet Psychiatry 9, 137–150. URL: https://www.thelancet.com/action/showFullText?pii=S2215036621003953 https://www.thelancet.com/action/showAbstract?pii=S2215036621003953 https://www.thelancet.com/journals/lanpsy/article/PIIS2215-0366(21)00395-3/abstract, doi:10.1016/S2215-0366(21)00395-3.

Flores-Gutiérrez, E., Cabrera-Muñoz, E.A., Vega-Rivera, N.M., Ortiz-López, L., Ramírez-Rodríguez, G.B., 2018. Exposure to Patterned Auditory Stimuli during Acute Stress Prevents Despair-Like Behavior in Adult Mice That Were Previously Housed in an Enriched Environment in Combination with Auditory Stimuli. Neural Plasticity 2018, 1–14. URL: https://www.hindawi.com/journals/np/2018/8205245/, doi:10.1155/2018/8205245.

Foldal, M.D., Blenkmann, A.O., Llorens, A., Knight, R.T., Solbakk, A.K., Endestad, T., 2020. The brain tracks auditory rhythm predictability independent of selective attention. Scientific Reports 2020 10:1 10, 7975–. URL: https://www.nature.com/articles/s41598-020-64758-y, doi:10.1038/s41598-020-64758-y.

Foo, Y.Z., O’Dea, R.E., Koricheva, J., Nakagawa, S., Lagisz, M., 2021. A practical guide to question formation, systematic searching and study screening for literature reviews in ecology and evolution. Methods in Ecology and Evolution 12, 1705–1720. URL: /doi/pdf/10.1111/2041-210X.13654 https://onlinelibrary.wiley.com/doi/abs/10.1111/2041-210X.13654 https://besjournals.onlinelibrary.wiley.com/doi/10.1111/2041-210X.13654, doi:10.1111/2041-210X.13654;PAGE:STRING:ARTICLE/CHAPTER.

Freitas Oliveira, B.R., Pires da Silva, T.W., Carvalho Alves, R.M., Soares Neto, J.R., Araújo Oliveira, C., Barbosa Guerreiro, D.F., Mazza Santos, M., Ramos da Costa, E., Picanço Diniz, C.W., Guerreiro Diniz, D., 2020. Classical Music and Environmental Enrichment Enhanced Spatial Memory and Learning and Increased Mouse Innate Tendency to Avoid Open Spaces. EC Neurology 13, 12–20. doi:10.31080/ecne.2021.13.00837.

Fu, Q., Qiu, R., Chen, L., Chen, Y., Qi, W., Cheng, Y., 2023. Music prevents stress-induced depression and anxiety-like behavior in mice. Translational Psychiatry 13, 317. URL: https://www.nature.com/articles/s41398-023-02606-z, doi:10.1038/s41398-023-02606-z.

Fu, Q., Qiu, R., Yao, T., Liu, L., Li, Y., Li, X., Qi, W., Chen, Y., Cheng, Y., 2025. Music therapy as a preventive intervention for postpartum depression: modulation of synaptic plasticity, oxidative stress, and inflammation in a mouse model. Translational Psychiatry 15, 143. URL: https://www.nature.com/articles/s41398-025-03370-y, doi:10.1038/s41398-025-03370-y.

Hedges, L., Gurevitch, J., Curtis, P., 1999. THE META-ANALYSIS OF RESPONSE RATIOS IN EXPERIMENTAL ECOLOGY. Ecology 80, 1150–1156. URL: https://esajournals.onlinelibrary.wiley.com/doi/abs/10.1890/0012-9658%281999%29080%5B1150%3ATMAORR%5D2.0.CO%3B2, doi:10.1890/0012-9658(1999)080[1150:TMAORR]2.0.CO;2.

Heilbron, M., Chait, M., 2018. Great Expectations: Is there Evidence for Predictive Coding in Auditory Cortex? Neuroscience 389, 54–73. URL: https://doi.org/10.1038/nn.2810, doi:10.1016/j.neuroscience.2017.07.061.

Holman, L., Head, M.L., Lanfear, R., Jennions, M.D., 2015. Evidence of Experimental Bias in the Life Sciences: Why We Need Blind Data Recording. PLOS Biology 13, e1002190. URL: https://journals.plos.org/plosbiology/article?id=10.1371/journal.pbio.1002190, doi:10.1371/journal.pbio.1002190.

Kelly, K.M., Mezuk, B., 2017. Predictors of remission from generalized anxiety disorder and major depressive disorder. Journal of Affective Disorders 208, 467–474. doi:10.1016/j.jad.2016.10.042.

Knight, P., Chellian, R., Wilson, R., Behnood-Rod, A., Panunzio, S., Bruijnzeel, A.W., 2021. Sex differences in the elevated plus-maze test and large open field test in adult Wistar rats. Pharmacology Biochemistry and Behavior 204, 173168. URL: https://doi.org/10.1371/journal.pone.0153327, doi:10.1016/j.pbb.2021.173168.

Koelsch, S., 2014. Brain correlates of music-evoked emotions. Nature Reviews Neuroscience 2014 15:3 15, 170–180. URL: https://www.nature.com/articles/nrn3666, doi:10.1038/nrn3666.

Krishnamurthy, T., Rao, B.V., 2025. Indian classical Mohana Raga (instrumental music) overcomes anxiety, depression and memory impairment in chronic unpredictable mild stress rat model – A behavioural study. Indian Journal of Traditional Knowledge URL: https://or.niscpr.res.in/index.php/IJTK/issue/view/386, doi:10.56042/ijtk.v24i4.12620.

Kühlmann, A.Y., de Rooij, A., Hunink, M.G., De Zeeuw, C.I., Jeekel, J., 2018. Music affects rodents: A systematic review of experimental research. Frontiers in Behavioral Neuroscience 12, 332220. URL: https://www.frontiersin.org, doi:10.3389/fnbeh.2018.00301.

Lagisz, M., Yang, Y., Young, S., Nakagawa, S., 2025. A practical guide to evaluating sensitivity of literature search strings for systematic reviews using relative recall. Research Synthesis Methods 16, 1–14. URL: https://www.cambridge.org/core/journals/research-synthesis-methods/article/practical-guide-to-evaluating-sensitivity-of-literature-search-strings-for-systematic-reviews-usinBC6A8387DAB7539D7F96EBD5965ECC32, doi:10.1017/RSM.2024.6.

Lajeunesse, M.J., 2011. On the meta-analysis of response ratios for studies with correlated and multi-group designs. Ecology 92, 2049–2055. URL: /doi/pdf/10.1890/11-0423.1 https://onlinelibrary.wiley.com/doi/abs/10.1890/11-0423.1 https://esajournals.onlinelibrary.wiley.com/doi/10.1890/11-0423.1, doi:10.1890/11-0423.1;WEBSITE:WEBSITE:ESAJOURNALS;REQUESTEDJOURNAL:JOURNAL:19399170;ISSUE:ISSUE:DOI.

Lazic, S.E., 2010. The problem of pseudoreplication in neuroscientific studies: is it affecting your analysis? BMC Neuroscience 2010 11:1 11, 5–. URL: https://link.springer.com/article/10.1186/1471-2202-11-5, doi:10.1186/1471-2202-11-5.

Le, J., Deng, W., Le, T., 2025. Music Therapy in Depression: Exploring Mechanisms and Efficacy in Rat Models. Brain Sciences 2025, Vol. 15, 15. URL: https://www.mdpi.com/2076-3425/15/4/338, doi:10.3390/brainsci15040338.

Leite-Almeida, H., Castelhano-Carlos, M.J., Sousa, N., 2022. New Horizons for Phenotyping Behavior in Rodents: The Example of Depressive-Like Behavior. Frontiers in Behavioral Neuroscience 15, 811987. URL: https://www.frontiersin.org, doi:10.3389/fnbeh.2021.811987.

Levitin, D.J., Tirovolas, A.K., 2009. Current advances in the cognitive neuroscience of music. Annals of the New York Academy of Sciences 1156, 211–231. URL: /doi/pdf/10.1111/j.1749-6632.2009.04417.x https://onlinelibrary.wiley.com/doi/abs/10.1111/j.1749-6632.2009.04417.x https://nyaspubs.onlinelibrary.wiley.com/doi/10.1111/j.1749-6632.2009.04417.x, doi:10.1111/j.1749-6632.2009.04417.x.

Li, W.J., Yu, H., Yang, J.M., Gao, J., Jiang, H., Feng, M., Zhao, Y.X., Chen, Z.Y., 2010. Anxiolytic effect of music exposure on BDNFMet/Met transgenic mice. Brain Research 1347, 71–79. doi:10.1016/j.brainres.2010.05.080.

Macartney, E.L., Lagisz, M., Nakagawa, S., 2022. The relative benefits of environmental enrichment on learning and memory are greater when stressed: A meta-analysis of interactions in rodents. Neuroscience & Biobehavioral Reviews 135, 104554. URL: https://www.sciencedirect.com/science/article/abs/pii/S0149763422000434?via%3Dihub, doi:10.1016/J.NEUBIOREV.2022.104554.

McDermott, J.H., Schultz, A.F., Undurraga, E.A., Godoy, R.A., 2016. Indifference to dissonance in native Amazonians reveals cultural variation in music perception. Nature 2016 535:7613 535, 547–550. URL: https://www.nature.com/articles/nature18635, doi:10.1038/nature18635.

Mcewen, B.S., Mirsky, A.E., Hatch, M.M., 2007. Physiology and neurobiology of stress and adaptation: central role of the brain. Physiological reviews 87, 873–904. URL: https://journals.physiology.org/doi/pdf/10.1152/physrev.00041.2006?download=true, doi:10.1152/physrev.00041.2006.

Mineur, Y.S., Belzung, C., Crusio, W.E., 2006. Effects of unpredictable chronic mild stress on anxiety and depression-like behavior in mice. Behavioural Brain Research 175, 43–50. URL: https://doi.org/10.1111/j.2042-7158.1988.tb05284.x, doi:10.1016/j.bbr.2006.07.029.

Mul, J.D., Mul, J.D., 2018. Voluntary exercise and depression-like behavior in rodents: are we running in the right direction? Journal of Molecular Endocrinology 60, R77–R95. URL: https://jme.bioscientifica.com/view/journals/jme/60/3/JME-17-0165.xml, doi:10.1530/JME-17-0165.

Nakagawa, S., Lagisz, M., Jennions, M.D., Koricheva, J., Noble, D.W., Parker, T.H., Sánchez-Tójar, A., Yang, Y., O’Dea, R.E., 2022. Methods for testing publication bias in ecological and evolutionary meta-analyses. Methods in Ecology and Evolution 13, 4–21. URL: /doi/pdf/10.1111/2041-210X.13724 https://onlinelibrary.wiley.com/doi/abs/10.1111/2041-210X.13724 https://besjournals.onlinelibrary.wiley.com/doi/10.1111/2041-210X.13724, doi:10.1111/2041-210X.13724;REQUESTEDJOURNAL:JOURNAL:2041210X;WEBSITE:WEBSITE:BESJOURNALS;WGROUP:STRING:PUBLICATION.

Nakagawa, S., Lagisz, M., O’Dea, R.E., Pottier, P., Rutkowska, J., Senior, A.M., Yang, Y., Noble, D.W., 2023a. orchaRd 2.0: An R package for visualising meta-analyses with orchard plots. Methods in Ecology and Evolution 14, 2003–2010. URL: /doi/pdf/10.1111/2041-210X.14152 https://onlinelibrary.wiley.com/doi/abs/10.1111/2041-210X.14152 https://besjournals.onlinelibrary.wiley.com/doi/10.1111/2041-210X.14152, doi:10.1111/2041-210X.14152;JOURNAL:JOURNAL:2041210X.

Nakagawa, S., Poulin, R., Mengersen, K., Reinhold, K., Engqvist, L., Lagisz, M., Senior, A.M., 2015. Metaanalysis of variation: Ecological and evolutionary applications and beyond. Methods in Ecology and Evolution 6, 143–152. URL: /doi/pdf/10.1111/2041-210X.12309 https://onlinelibrary.wiley.com/doi/abs/10.1111/2041-210X.12309 https://besjournals.onlinelibrary.wiley.com/doi/10.1111/2041-210X.12309, doi:10.1111/2041-210X.12309;WEBSITE:WEBSITE:BESJOURNALS;WGROUP:STRING:PUBLICATION.

Nakagawa, S., Santos, E.S., 2012. Methodological issues and advances in biological meta-analysis. Evolutionary Ecology 2012 26:5 26, 1253–1274. URL: https://link.springer.com/article/10.1007/s10682-012-9555-5, doi:10.1007/s10682-012-9555-5.

Nakagawa, S., Yang, Y., Macartney, E.L., Spake, R., Lagisz, M., 2023b. Quantitative evidence synthesis: a practical guide on meta-analysis, meta-regression, and publication bias tests for environmental sciences. Environmental Evidence 12, 8. URL: https://pmc.ncbi.nlm.nih.gov/articles/PMC11378872/, doi:10.1186/s13750-023-00301-6.

Nestler, E.J., Barrot, M., DiLeone, R.J., Eisch, A.J., Gold, S.J., Monteggia, L.M., 2002. Neurobiology of Depression. Neuron 34, 13–25. URL: https://doi.org/10.1677/joe.0.1600001, doi:10.1016/S0896-6273(02)00653-0.

Noble, D.W., Lagisz, M., O’dea, R.E., Nakagawa, S., 2017. Nonindependence and sensitivity analyses in ecological and evolutionary meta-analyses. Molecular Ecology 26, 2410–2425. URL: /doi/pdf/10.1111/mec.14031 https://onlinelibrary.wiley.com/doi/abs/10.1111/mec.14031 https://onlinelibrary.wiley.com/doi/10.1111/mec.14031, doi:10.1111/MEC.14031;WEBSITE:WEBSITE:PERICLES;ISSUE:ISSUE:DOI.

O’Dea, R.E., Lagisz, M., Jennions, M.D., Koricheva, J., Noble, D.W., Parker, T.H., Gurevitch, J., Page, M.J., Stewart, G., Moher, D., Nakagawa, S., 2021. Preferred reporting items for systematic reviews and meta-analyses in ecology and evolutionary biology: a PRISMA extension. Biological Reviews 96, 1695–1722. URL: /doi/pdf/10.1111/brv.12721 https://onlinelibrary.wiley.com/doi/abs/10.1111/brv.12721 https://onlinelibrary.wiley.com/doi/10.1111/brv.12721, doi:10.1111/brv.12721.

Ortega, S., Lenz, A., Lundgren, E., Mizuno, A., Poo-Hernandez, S., Nakagawa, S., Lagisz, M., 2026. Music exposure reduces anxiety- and depression-like behavior in rodents: a systematic review and multilevel meta-analysis. URL: https://santiago-0rtega.github.io/Music_on_anxiety_depression_meta_analysis/, doi:10.5281/zenodo.18792126.

Ouzzani, M., Hammady, H., Fedorowicz, Z., Elmagarmid, A., 2016. Rayyan—a web and mobile app for systematic reviews. Systematic Reviews 2016 5:1 5, 210–. URL: https://link.springer.com/article/10.1186/s13643-016-0384-4, doi:10.1186/S13643-016-0384-4.

Page, M.J., McKenzie, J.E., Bossuyt, P.M., Boutron, I., Hoffmann, T.C., Mulrow, C.D., Shamseer, L., Tetzlaff, J.M., Akl, E.A., Brennan, S.E., Chou, R., Glanville, J., Grimshaw, J.M., Hróbjartsson, A., Lalu, M.M., Li, T., Loder, E.W., Mayo-Wilson, E., McDonald, S., McGuinness, L.A., Stewart, L.A., Thomas, J., Tricco, A.C., Welch, V.A., Whiting, P., Moher, D., 2021. The PRISMA 2020 statement: an updated guideline for reporting systematic reviews. BMJ 372. URL: https://www.bmj.com/content/372/bmj.n71 https://www.bmj.com/content/372/bmj.n71.abstract, doi:10.1136/bmj.n71.

Pangemanan, L., Irwanto Maramis, M.M., 2024. Mozart K488 Addition Can Improve Depressive-Like Behavior in Rats: In Search of Better Management. Pharmacognosy Journal 16, 348–354. doi:10.5530/pj.2024.16.53.

Papadakakis, A., Sidiropoulou, K., Panagis, G., 2019. Music exposure attenuates anxiety- and depression-like behaviors and increases hippocampal spine density in male rats. Behavioural Brain Research 372, 112023. URL: https://linkinghub.elsevier.com/retrieve/pii/S0166432819303249, doi:10.1016/j.bbr.2019.112023.

Pendersen, T., 2025. patchwork: The Composer of Plots. URL: https://patchwork.data-imaginist.com/.

Pick, J.L., Nakagawa, S., Noble, D.W., 2019. Reproducible, flexible and high-throughput data extraction from primary literature: The metaDigitise r package. Methods in Ecology and Evolution 10, 426–431. URL: /doi/pdf/10.1111/2041-210X.13118 https://onlinelibrary.wiley.com/doi/abs/10.1111/2041-210X.13118 https://besjournals.onlinelibrary.wiley.com/doi/10.1111/2041-210X.13118, doi:10.1111/2041-210X.13118;REQUESTEDJOURNAL:JOURNAL:2041210X;WEBSITE:WEBSITE:BESJOURNALS;WGROUP:STRING:PUBLICATION.

Posit team, 2026. RStudio: Integrated Development Environment for R. URL: http://www.posit.co/.

R Core Team, 2025. R: A Language and Environment for Statistical Computing. URL: https://www.R-project.org/.

Ren, J., Lu, J., 2024. Heavy metal Music, Hip-hop Music and Construction Noise Induces Depressive Symptoms in mice. ASEAN Journal of Psychiatry 25, 1–14.

Rickard, N.S., Toukhsati, S.R., Field, S.E., 2005. The effect of music on cognitive performance: Insight from neurobiological and animal studies. Behavioral and Cognitive Neuroscience Reviews 4, 235–261. URL: /doi/pdf/10.1177/1534582305285869?download=true, doi:10.1177/1534582305285869.

Ridley, M., Rao, G., Schilbach, F., Patel, V., 2020. Poverty, depression, and anxiety: Causal evidence and mechanisms. Science 370. URL: /doi/pdf/10.1126/science.aay0214?download=true, doi:10.1126/science.aay0214.

Rizzolo, L., Leger, M., Corvaisier, S., Groussard, M., Platel, H., Bouet, V., Schumann-Bard, P., Freret, T., 2021. Long-Term Music Exposure Prevents Age-Related Cognitive Deficits in Rats Independently of Hippocampal Neurogenesis. Cerebral Cortex 31, 620–634. URL: https://academic.oup.com/cercor/article/31/1/620/5909651, doi:10.1093/cercor/bhaa247.

Russo, S.J., Murrough, J.W., Han, M.H., Charney, D.S., Nestler, E.J., 2012. Neurobiology of resilience. Nature Neuroscience 15, 1475–1484. URL: https://www.nature.com/articles/nn.3234, doi:10.1038/nn.3234.

Saghari, H., Sheibani, V., Esmaeilpour, K., ur Rehman, N., 2021. Music Alleviates Learning and Memory Impairments in an Animal Model of Post-Traumatic Stress Disorder. Biointerface Research in Applied Chemistry 11, 7775–7784. URL: https://biointerfaceresearch.com/wp-content/uploads/2020/07/20695837111.77757784.pdf, doi:10.33263/BRIAC111.77757784.

Sampaio, W.C.M., Ribeiro, M.C., Costa, L.F., Souza, W.C.d., Castilho, G.M.d., Assis, M.S.d., Carneiro, F.P., Marchiori, D., Lima, N.T.d., Cavalcanti, P.P., Ferreira, V.M., 2017. Effect of music therapy on the developing central nervous system of rats. Psychology & Neuroscience 10, 176–188. URL: https://doi.apa.org/doi/10.1037/pne0000087, doi:10.1037/pne0000087.

Saré, R.M., Lemons, A., Smith, C.B., 2021. Behavior Testing in Rodents: Highlighting Potential Confounds Affecting Variability and Reproducibility. Brain Sciences 2021, Vol. 11, 11. URL: https://www.mdpi.com/2076-3425/11/4/522, doi:10.3390/brainsci11040522.

Senior, A.M., Viechtbauer, W., Nakagawa, S., 2020. Revisiting and expanding the meta-analysis of variation: The log coefficient of variation ratio. Research Synthesis Methods 11, 553–567. URL: /doi/pdf/10.1002/jrsm.1423 https://onlinelibrary.wiley.com/doi/abs/10.1002/jrsm.1423 https://onlinelibrary.wiley.com/doi/10.1002/jrsm.1423, doi:10.1002/JRSM.1423;WGROUP:STRING:PUBLICATION.

Sih, A., Mathot, K.J., Moirón, M., Montiglio, P.O., Wolf, M., Dingemanse, N.J., 2015. Animal personality and state-behaviour feedbacks: A review and guide for empiricists. Trends in Ecology and Evolution 30, 50–60. URL: https://pubmed.ncbi.nlm.nih.gov/25498413/, doi:10.1016/j.tree.2014.11.004.

Simpson, J., Kelly, J.P., 2011. The impact of environmental enrichment in laboratory rats—Behavioural and neurochemical aspects. Behavioural Brain Research 222, 246–264. URL: https://doi.org/10.1093/ilar.46.2.83, doi:10.1016/j.bbr.2011.04.002.

Slattery, D.A., Cryan, J.F., 2012. Using the rat forced swim test to assess antidepressant-like activity in rodents. Nature protocols 7, 1009–1014. URL: https://pubmed.ncbi.nlm.nih.gov/22555240/, doi:10.1038/nprot.2012.044.

Smith, S.M., Vale, W.W., 2006. The role of the hypothalamic-pituitary-adrenal axis in neuroendocrine responses to stress. Dialogues in Clinical Neuroscience 8, 383. URL: https://pmc.ncbi.nlm.nih.gov/articles/PMC3181830/, doi:10.31887/dcns.2006.8.4/ssmith.

Thoma, M.V., La Marca, R., Brönnimann, R., Finkel, L., Ehlert, U., Nater, U.M., 2013. The Effect of Music on the Human Stress Response. PLOS ONE 8, e70156. URL: https://journals.plos.org/plosone/article?id=10.1371/journal.pone.0070156, doi:10.1371/journal.pone.0070156.

Uşak, T., Dal, A., Yanık, H., Elibol, B., 2020. Effects of music on stress induced hormones and oxidative stress levels. Cukurova Medical Journal 45, 1493–1498. URL: http://dergipark.org.tr/en/doi/10.17826/cumj.735738, doi:10.17826/cumj.735738.

Viechtbauer, W., 2010. Conducting Meta-Analyses in R with the metafor Package. Journal of Statistical Software 36, 1–48. URL: https://www.jstatsoft.org/index.php/jss/article/view/v036i03, doi:10.18637/JSS.V036.I03.

Voelkl, B., Altman, N.S., Forsman, A., Forstmeier, W., Gurevitch, J., Jaric, I., Karp, N.A., Kas, M.J., Schielzeth, H., Van de Casteele, T., Würbel, H., 2020. Reproducibility of animal research in light of biological variation. Nature reviews. Neuroscience 21, 384–393. URL: https://pubmed.ncbi.nlm.nih.gov/32488205/, doi:10.1038/s41583-020-0313-3.

Walf, A.A., Frye, C.A., 2007. The use of the elevated plus maze as an assay of anxiety-related behavior in rodents. Nature Protocols 2007 2:2 2, 322–328. URL: https://www.nature.com/articles/nprot.2007.44, doi:10.1038/nprot.2007.44.

Westneat, D.F., Wright, J., Dingemanse, N.J., 2015. The biology hidden inside residual within-individual phenotypic variation. Biological Reviews 90, 729–743. URL: /doi/pdf/10.1111/brv.12131 https://onlinelibrary.wiley.com/doi/abs/10.1111/brv.12131 https://onlinelibrary.wiley.com/doi/10.1111/brv.12131, doi:10.1111/brv.12131.

Wickham, H., 2011. ggplot2. WIREs Computational Statistics 3, 180–185. URL: https://wires.onlinelibrary.wiley.com/doi/10.1002/wics.147, doi:10.1002/wics.147.

Wickham, H., Averick, M., Bryan, J., Chang, W., D’, L., Mcgowan, A., François, R., Grolemund, G., Hayes, A., Henry, L., Hester, J., Kuhn, M., Lin Pedersen, T., Miller, E., Bache, S.M., Müller, K., Ooms, J., Robinson, D., Seidel, D.P., Spinu, V., Takahashi, K., Vaughan, D., Wilke, C., Woo, K., Yutani, H., 2019. Welcome to the Tidyverse. Journal of Open Source Software 4, 1686. URL: https://joss.theoj.org/papers/10.21105/joss.01686, doi:10.21105/JOSS.01686.

Willner, P., 2017. The chronic mild stress (CMS) model of depression: History, evaluation and usage. Neurobiology of Stress 6, 78–93. URL: https://doi.org/10.1038/mp.2008.106, doi:10.1016/j.ynstr.2016.08.002.

de Witte, M., Pinho, A.d.S., Stams, G.J., Moonen, X., Bos, A.E., van Hooren, S., 2022. Music therapy for stress reduction: a systematic review and meta-analysis. Health Psychology Review 16, 134–159. URL: https://www.tandfonline.com/doi/pdf/10.1080/17437199.2020.1846580, doi:10.1080/17437199.2020.1846580.

Würbel, H., 2001. Ideal homes? Housing effects on rodent brain and behaviour. Trends in Neurosciences 24, 207–211. URL: https://doi.org/10.1038/35044558, doi:10.1016/S0166-2236(00)01718-5.

Yang, Y., Zhu, Y., Chen, B., Li, Y., Li, H., Zhang, G., 2025. Effects of Physical Exercise on Depression and Anxiety-Like Behaviors in Rodent Models of Maternal Separation: A Systematic Review and Meta-Analysis. Journal of Sport for All and Recreation 7, 777–787. URL: https://dergipark.org.tr/en/pub/jsar/article/1720636, doi:10.56639/jsar.1720636.

