## Supplementary Material for "Music exposure reduces anxiety- and depression-like behavior in rodents: a systematic review and multilevel meta-analysis"

February 27, 2026

### S1 Literature searches

#### S1.1 Web of Science Core Collection search string (2025/09/24; 750 hits):

(TS=((music* OR Mozart OR “auditory enrich*”OR “acoustic enrich*”) AND (depress* OR anxi* OR distress* OR stress* OR despair* OR CUMS OR CMS or anhedonia OR PTSD OR PPD OR “emotional state*”OR “open field”OR “elevated plus maze”OR “forced swim*”OR “sucrose preference”OR “light dark box”OR “marble burying”OR “Novelty Suppressed Feeding”OR “tail suspension”OR behav* OR “Maternal Separat*”OR “Social Defeat “OR “Learned Helplessness”) AND (mice OR mouse OR rat OR rats OR rodent* OR animal* OR “non human”OR “pre clinical”OR murine OR “Swiss Webster”OR “Sprague-Dawley”OR “Wistar”OR “Flinders Sensitive Line”OR “Holtzman Albino” OR “Long Evans”OR “C57BL/6*”OR “BALB/c”OR “ICR”OR “DBA/2”OR “FVB/N”)))

#### S1.2 Scopus search string (2025/09/24; 1301 hits):

TITLE-ABS-KEY((music* OR Mozart OR “auditory enrich*”OR “acoustic enrich*”) AND (depress* OR anxi* OR distress* OR stress* OR despair* OR CUMS OR CMS OR anhedonia OR PTSD OR PPD OR “emotional state*” OR “open field”OR “elevated plus maze”OR “forced swim*”OR “sucrose preference”OR “light dark box”OR “marble burying”OR “Novelty Suppressed Feeding”OR “tail suspension”OR behav* OR “Maternal Separat*”OR “Social Defeat”OR “Learned Helplessness”) AND (mice OR mouse OR rat OR rats OR rodent* OR animal* OR “non human” OR “pre clinical”OR murine OR “Swiss Webster”OR “Sprague-Dawley”OR “Wistar”OR “Flinders Sensitive Line”OR “Holtzman Albino”OR “Long Evans”OR “C57BL/6*”OR “BALB/c”OR “ICR”OR “DBA/2”OR “FVB/N”))

#### S1.3 PubMed search string (2025/09/24; 377 hits):

(music*[Title/Abstract] OR Mozart[Title/Abstract] OR “auditory enrich*”[Title/Abstract] OR “acoustic enrich*”[Title/Abstract]) AND (depress*[Title/Abstract] OR anxi*[Title/Abstract] OR distress*[Title/Abstract] OR stress*[Title/Abstract] OR despair*[Title/Abstract] OR CUMS[Title/Abstract] OR CMS[Title/Abstract] OR anhedonia[Title/Abstract] OR PTSD[Title/Abstract] OR PPD[Title/Abstract] OR “emotional state*”[Title/Abstract] OR “open field”[Title/Abstract] OR “elevated plus maze” [Title/Abstract] OR “forced swim*”[Title/Abstract] OR “sucrose preference”[Title/Abstract] OR “light dark box”[Title/Abstract] OR “marble burying”[Title/Abstract] OR “Novelty Suppressed Feeding”[Title/Abstract] OR “tail suspension”[Title/Abstract] OR behav*[Title/Abstract] OR “Maternal Separat*”[Title/Abstract] OR “Social Defeat”[Title/Abstract] OR “Learned Helplessness”[Title/Abstract]) AND (mice[Title/Abstract] OR mouse[Title/Abstract] OR rat[Title/Abstract] OR rats[Title/Abstract] OR rodent*[Title/Abstract] OR animal*[Title/Abstract] OR “non human” [Title/Abstract] OR “pre clinical”[Title/Abstract] OR murine[Title/Abstract] OR “Swiss Webster”[Title/Abstract] OR “Sprague-Dawley”[Title/Abstract] OR “Wistar”[Title/Abstract] OR “Flinders Sensitive Line”[Title/Abstract] OR “Holtzman Albino”[Title/Abstract] OR “Long Evans”[Title/Abstract] OR “C57BL/6*”[Title/Abstract] OR “BALB/c”[Title/Abstract] OR “ICR”[Title/Abstract] OR “DBA/2”[Title/Abstract] OR “FVB/N”[Title/Abstract])

##### S1.4 OpenAlex search string (2025/09/29; 1126):

(music OR Mozart OR ”auditory enrich*” OR ”acoustic enrich*”) AND (depress* OR anxi* OR distress* OR stress* OR despair* OR CUMS OR CMS OR anhedonia OR PTSD OR PPD OR ”emotional state” OR ”open field” OR ”elevated plus maze” OR ”forced swim” OR ”sucrose preference” OR ”light dark box” OR ”marble burying” OR ”Novelty Suppressed Feeding” OR ”tail suspension” OR behav* OR ”Maternal Separat*” OR ”Social Defeat” OR ”Learned Helplessness”) AND (mice OR mouse OR rat* OR rodent* OR animal* OR ”non human” OR preclinical OR murine OR ”Swiss Webster” OR ”Sprague-Dawley” OR Wistar OR ”Flinders Sensitive Line” OR ”Holtzman Albino” OR ”Long Evans” OR C57BL/6 OR BALB/c OR ICR OR DBA/2 OR FVB/N)

##### S1.5 Supplementary searches

<table>
<tr><th>Language</th><th>Search Query</th></tr>
<tr><td>English</td><td>(music OR Mozart) AND (depress OR anxiety OR “open field”OR “elevated plus maze”OR “forced swim” OR “light dark box”OR “marble bury”OR “novelty suppressed feed”OR “tail suspension”) AND (mice OR mouse OR rat OR rodent)</td></tr>
<tr><td>Spanish</td><td>(música|Mozart) AND (depresión|ansiedad|”campo abierto”|”laberinto en cruz elevado”|”nado forzado”| “caja de luz-oscuridad”|”enterramiento de canicas”|”alimentación suprimida por la novedad”|”suspensión de la cola”) AND (ratones|ratón|rata|roedor)</td></tr>
<tr><td>Japanese</td><td>(音楽 | モーツァルト) AND (うつ病 | 不安 | オープンフィールド | 高架式十字迷路 | 強制水泳 | 明暗箱 | 新規環境摂食抑制試験 | 尾懸垂 | 鬱病 | ガラス玉覆い隠し | ビー玉覆い隠し) AND (マウス | ラット | げっ歯類 | 齧歯類)</td></tr>
<tr><td>Polish</td><td>(muzyka|Mozart) AND (depresja|lęk|”otwarte pole”|”podwyższony labirynt krzyżowy”|”wymuszony test pływania”|”test światło-ciemność”|”test zakopywania kulek”|”hamowanie jedzenia w nowym otoczeniu”|” test zawieszenia za ogon”) AND (myszy|mysz|szczur|gryzoń)</td></tr>
<tr><td>Russian</td><td>(музыка|Моцарт) AND (депрессия|тревожносты|”открытое поле”|”приподнятый крестообразный лабиринт”|”принудительное плавание”| тест свет-темнота”|”закапывание шариков”|” подавление потребления пищи в новой обстановке”|”подвешивание за хвост”) AND (мыши|мышь|крыса|грызун)</td></tr>
</table>

Search strategies adapted for five languages.

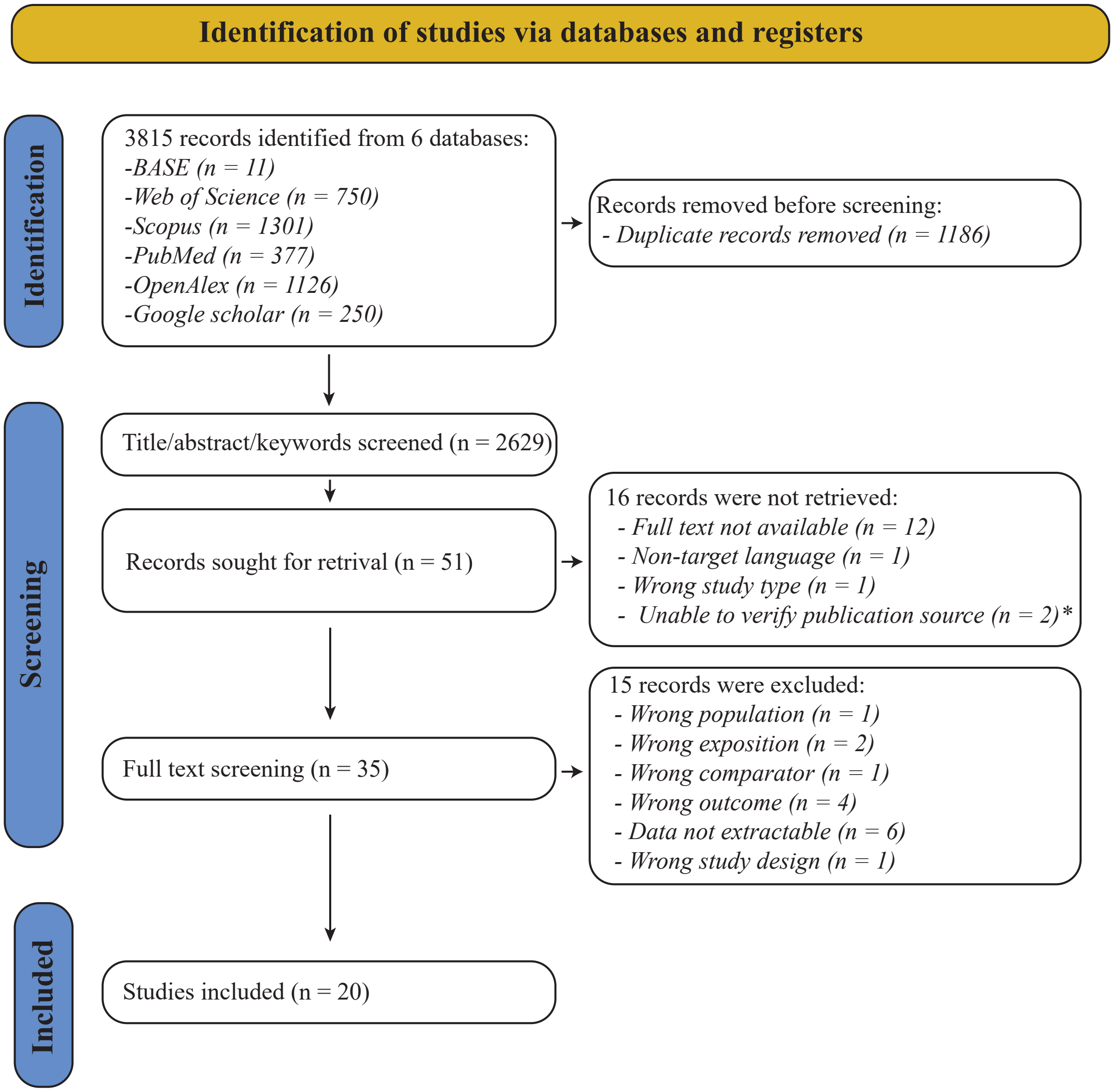

* During the screening process, two records retrieved from Google Scholar presented non-existent or conflicting bibliographic information. Despite attempts to verify the original publication source, the correct citation could not be determined

Figure S1: PRISMA flowchart

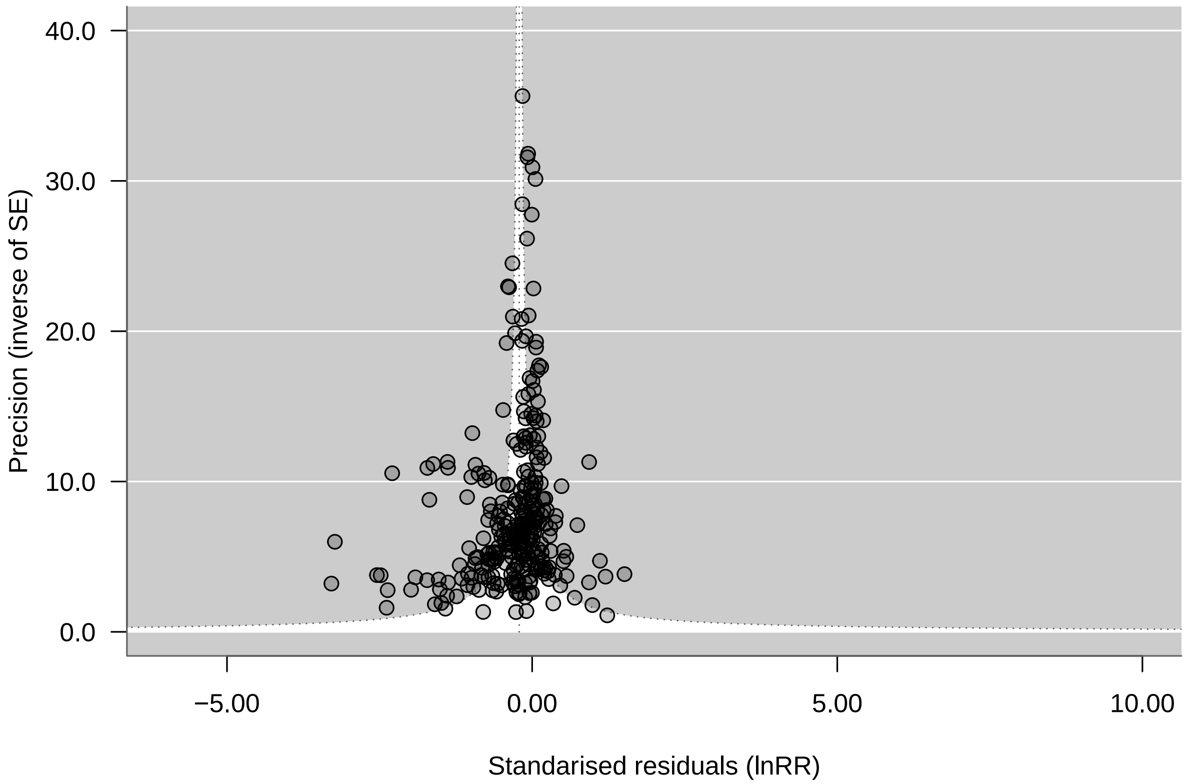

Figure S2: Funnel plot of effect sizes against inverse standard error

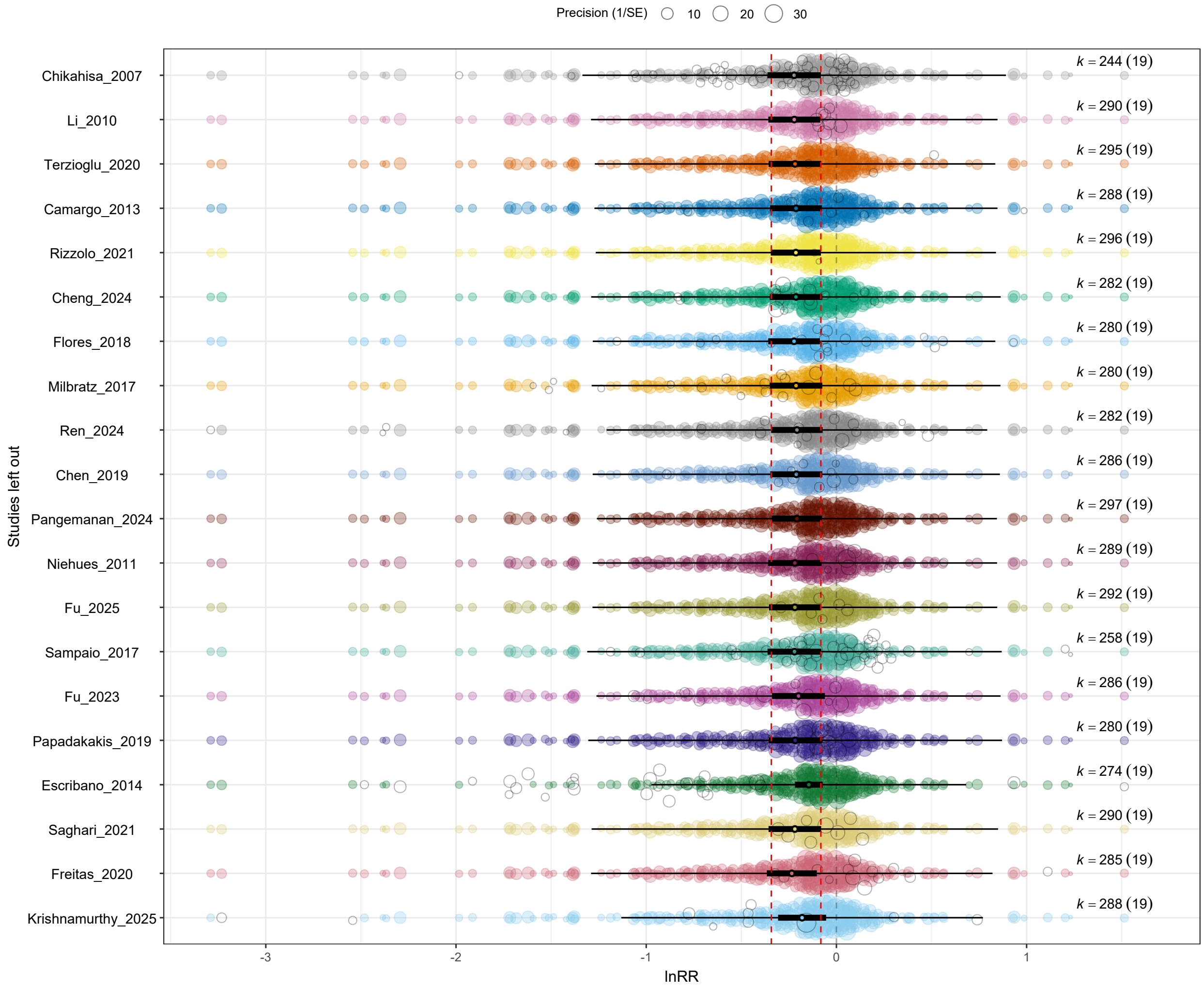

Figure S3: Leave-one-out sensitivity analysis of mean behavioral effects (ln $RR$). Each point represents the pooled effect size estimate obtained after sequentially excluding one study from the dataset and refitting the multilevel meta-analytic model. Horizontal lines indicate 95% confidence intervals around each leave-one-out estimate. The vertical dashed line represents the overall pooled estimate based on the full dataset. Consistency of the estimates and overlapping confidence intervals indicate that no single study exerted disproportionate influence on the overall effect of music exposure on anxiety- and depression-like behaviors.

#### S2 Tables

Table S1: Definitions and coding rules for moderators used in meta-regressions.

| Moderator | Definition and coding |
|---|---|
| **LifeStage_Exposure** | Age category at music exposure: Neonatal (birth–∼1 week), Infantile (∼1–2 weeks), Juvenile (∼2–4 weeks), Adolescent/Peripubertal (∼4–8 weeks), Young adult (∼2–5 months), Adult (∼5–18 months), or Old (>18 months). Multiple ages coded as Mixed. Missing or unclear information coded as Not reported or Unclear. |
| **Experimental_Design** | Design used to estimate music effects: Posttest-only (independent groups measured once), Repeated-measures (same animals measured across time or conditions), or Factorial (music crossed with additional factors). |
| **Behavioral_Mechanism** | Indicates whether behavior was measured under baseline conditions (Innate) or following an induction procedure intended to elevate anxiety/depression-like phenotypes (Induced; e.g., chronic stress, maternal separation). |
| **Experimental_Procedures** | Indicates whether a sham procedure controlled for surgical or pharmacological manipulation. Coded as None or Sham. |
| **Relative_Timing** | Timing of music exposure relative to testing: Before (completed prior to testing), Concurrent (during testing), Both (before and during), or Not specified. |
| **Music_Exposure_Duration** | Total exposure categorized as Acute (1–7 days), Short-term (8–28 days), Medium-term (29–60 days), or Long-term (>60 days). |
| **Meta_Genre** | Broad classification of stimulus type: Western art/orchestral, Popular contemporary (post-1950 mainstream styles), Traditional/folk/world, Mixed (multiple genres), or Unclear. |
| **Control_Condition** | Condition experienced by control animals during exposure: Silence, Ambient sound, or White noise. |
| **Outcome_Type** | Behavioral construct measured, coded as Anxiety, Depression, Both, or Unclear if not specified. |
| **Assay_Type** | Specific behavioral assay (e.g., Elevated Plus Maze, Open Field Test, Light–Dark Box, Forced Swim Test, Tail Suspension Test, Sucrose Preference Test, Novelty-Suppressed Feeding). |
| **Higher_Better** | Directionality indicator for harmonizing effect sizes. Yes = higher mean reflects lower anxiety/depression; No = higher mean reflects higher anxiety/depression. |

Table S2: Reporting transparency and risk-of-bias indicators across included studies. Values show the proportion of studies reporting each item.

| **Item** | **Proportion (%)** |
|---|---|
| Stimulus reporting | 55% |
| Control stimulus reporting | 10% |
| Timing reporting | 25% |
| Randomization reported | 65% |
| Blinding reported | 35% |
| Attrition handling reported | 10% |
| Selective reporting flagged | 20% |
| Use of formal RoB guideline | 0% |

Table S3: Included studies.

| Title | Year | Authors | Journal | DOI |
|---|---|---|---|---|
| Exposure to Patterned Auditory Stimuli during Acute Stress Prevents Despair-Like Behavior in Adult Mice That Were Previously Housed in an Enriched Environment in Combination with Auditory Stimuli | 2018 | Flores-Gutiérrez et al. | Neural Plasticity | 10.1155/2018/8205245 |
| Music therapy as a preventive intervention for postpartum depression: modulation of synaptic plasticity, oxidative stress, and inflammation in a mouse model | 2025 | Fu et al. | Translational Psychiatry | 10.1038/s41398-025-03370-y |
| Effect of music therapy on the developing central nervous system of rats | 2017 | Sampaio et al. | Psychology and Neuroscience | 10.1037/pne0000087 |
| Anxiolytic effect of music depends on ovarian steroid in female mice | 2007 | Chikahisa et al. | Behavioural Brain Research | 10.1016/j.bbr.2007.01.010 |
| Music prevents stress-induced depression and anxiety-like behavior in mice | 2023 | Fu et al. | Translational Psychiatry | 10.1038/s41398-023-02606-z |
| Music exposure attenuates anxiety- and depression-like behaviors and increases hippocampal spine density in male rats | 2019 | Papadakakis et al. | Behavioural Brain Research | 10.1016/j.bbr.2019.112023 |
| Role of noise and music as anxiety modulators: Relationship with ovarian hormones in the rat | 2014 | Escribano et al. | Applied Animal Behaviour Science | 10.1016/j.applanim.2013.12.006 |
| Music Alleviates Learning and Memory Impairments in an Animal Model of Post-Traumatic Stress Disorder | 2021 | Saghari et al. | Biointerface Research in Applied Chemistry | 10.33263/BRIAC111.77757784 |
| Indian classical Mohana Raga (instrumental music) overcomes anxiety, depression and memory impairment in chronic unpredictable mild stress rat model –A behavioural study | 2025 | Krishnamurthy and Venkanna Rao | Indian Journal of Traditional Knowledge | 10.56042/ijtk.v24i4.12620 |
| Adjuvant effects of classical music on simvastatin induced reduction of anxiety but not object recognition memory in rats | 2013 | Camargo et al. | Psychology and Neuroscience | 10.3922/j.psns.2013.3.19 |
| Anxiolytic effect of music exposure on BDNFMet/Met transgenic mice | 2010 | Li et al. | Brain Research | 10.1016/j.brainres.2010.05.080 |
| Long-Term Music Exposure Prevents Age-Related Cognitive Deficits in Rats Independently of Hippocampal Neurogenesis | 2021 | Rizzolo et al. | Cerebral Cortex | 10.1093/cercor/bhaa247 |
| Effects of music on stress induced hormones and oxidative stress levels | 2020 | Uşak et al. | Cukurova Medical Journal | 10.17826/cumj.735738 |
| Classical Music and Environmental Enrichment Enhanced Spatial Memory and Learning and Increased Mouse Innate Tendency to Avoid Open Spaces | 2020 | Freitas Oliveira et al. | EC Neurology | 10.31080/ecne.2021.13.00837 |

| Title | Year | Authors | Journal | DOI |
|---|---|---|---|---|
| Light and classical music therapies attenuate chronic unpredictable mild stress-induced depression via BDNF signaling pathway in mice | 2024 | Cheng et al. | Heliyon | 10.1016/j.heliyon.2024.e34196 |
| Cocoa and classical music: effect on anxiety and antioxidant activity in Wistar rats | 2017 | Milbratz de Camargo et al. | Archivos Latinoamericanos de Nutrición | NA |
| Heavy metal Music, Hip-hop Music and Construction Noise Induces Depressive Symptoms in mice | 2024 | Ren and Lu | ASEAN Journal of Psychiatry | NA |
| Regular Music Exposure in Juvenile Rats Facilitates Conditioned Fear Extinction and Reduces Anxiety after Foot Shock in Adulthood | 2019 | Chen et al. | BioMed Research International | 10.1155/2019/8740674 |
| Mozart K488 Addition Can Improve Depressive-Like Behavior in Rats: In Search of Better Management | 2024 | Pangemanan et al. | Pharmacognosy Journal | 10.5530/pj.2024.16.53 |
| The Power of Classic Music to Reduce Anxiety in Rats Treated with Simvastatin | 2011 | Niehues da Cruz et al. | Basic and Clinical Neuroscience | NA |

Table S4: Excluded studies.

| Title | Year | Authors | Journal | DOI | Reason for exclusion |
|---|---|---|---|---|---|
| The effect of music exposure in juvenile stage on anxiety-like behavior and fear extinction in adult rat | 2012 | Liang et al. | Chinese Journal of Behavioral Medicine and Brain Science | 10.3760/CMA.J.ISSN.1674-6554.2012.04.008 | Full-text unavailable/Not in target language |
| Investigation of the Influence of Music on Behavioral Findings by Valproic Acid-Induced Autism Model in Rats | 2018 | Tümentemur et al. | Turkish Society of Physiological Sciences 44th National Physiology Congress | NA | Wrong type of study/Unable to verify publication source |
| Dopamine dynamics in chronic pain: music-induced, sex-dependent, behavioral effects in mice | 2024 | Flores-García et al. | Pain Reports | 10.1097/PR9.0000000000001205 | Wrong comparator |
| [Effect of electroacupuncture intervention on behavioral changes and hippocampal excitatory amino acid transporter mRNA expression in depression rats] | 2013 | Ji et al. | Zhen Ci Yan Jiu | 10.13702/j.1000-0607.2013.03.006 | Full-text unavailable/Not in target language |
| Vliyanie razlichnykh zvukovykh stressorov na povedenie laboratornykh myshei v teste ”otkrytoe pole” | 2016 | Germanovich | Aktual'nye problemy biologicheskoi i khimicheskoi ekologii | NA | Data not extractable |
| Study of the effect of Kalyani raga in Anxiety-like conditions in female Wistar rats | 2024 | Rathod and Vaidya | International Journal of Ayurvedic Medicine | 10.47552/ijam.v15i1.2977 | Wrong study design/Wrong outcome |
| ROLE OF HO-1 IN THE EFFECTOR PHASE OF ARTHRITIS FINASTERIDE AND ALLOPREGENENOLONE ADMINISTRATION ON THE ANXIOLYTIC EFFECT OF MUSIC IN FEMALE RATS. INTRODUCTION | 2011 | Lopez et al. | BASIC and CLINICAL PHARMACOLOGY and TOXICOLOGY | NA | Unable to verify publication source |
| Rapid auditory processing and MGN morphology in microgyric rats reared in varied acoustic environments | 2002 | Peiffer et al. | Developmental Brain Research | 10.1016/S0165-3806(02)00472-8 | Wrong outcome |
| Mozart KV 448 Menurunkan Densitas dan Aktivitas Neuroglia Hipokampus Mencit (Mus musculus) Selama Stres Prenatal No. 416-KE | 2017 | Kusumarini et al. | Jurnal Sain Veteriner | 10.22146/jsv.29279 | Full-text unavailable/Not in target language |
| Patterns of movement in the open field of the rat with autistic behavior subjected to musical stimulation | 2018 | Monje-Reyna et al. | Neurobiología | NA | Data not extractable |
| Musicotherapy treating rat models with post-stroke depression by reducing expressions of serum IL-10 and hippocampal PDE4A | 2019 | Lin et al. | BASIC and CLINICAL PHARMACOLOGY and TOXICOLOGY | NA | Unable to verify publication source/ |
| Music and methamphetamine: Conditioned cue-induced increases in locomotor activity and dopamine release in rats | 2011 | Polston et al. | Pharmacology Biochemistry and Behavior | 10.1016/j.pbb.2010.11.024 | Wrong outcome |
| Mouse Ethology: Effect of different human and rodentized music upon the social and individual behaviour, general feeling and genetics-environment interaction of mice II. Effect of original and five octaves higher music of Bach and MOZART on the behaviour of mice of different genotypes | 2016 | Korsós et al. | Magyar Állatorvosok Lapja | NA | Full-text unavailable/Not in target language |

| Title | Year | Authors | Journal | DOI | Reason for exclusion |
|---|---|---|---|---|---|
| Molecular mechanism of reward treatment ameliorating chronic stress-induced depressive-like behavior assessed by sequencing miRNA and mRNA in medial prefrontal cortex | 2020 | An et al. | Biochemical and Biophysical Research Communications | 10.1016/j.bbrc.2020.05.158 | Wrong exposure |
| Is Mice Behavior Influenced by Music? | 2013 | Falkenhorst | Szent István University | NA | Full-text unavailable/Not in target language |
| Influence of different environmental effects of human origin (socialization, music, noisemusic, noise) upon the rats' behaviour. Part 3. Effect of different noises on the open-field test behaviour | 2013 | Fekete et al. | Magyar Állatorvosok Lapja | NA | Full-text unavailable/Not in target language |
| Influence of different environmental effects of human origin (socialization, music, noisemusic, noise) upon the rats' behaviour. Part 2. Do rats react on human music? | 2013 | Fekete and Bernitsa | Magyar Állatorvosok Lapja | NA | Full-text unavailable/Not in target language |
| Influence of different environmental effects of human origin (socialization, music, noisemusic, noise) upon the rats' behaviour. Part 1 | 2013 | Korsós and Fekete | Magyar Állatorvosok Lapja | NA | Full-text unavailable/Not in target language |
| How the Different Noise Types May Influence the Open-field Behavior of Rats? | 2013 | Sukikara | Department of Animal Breeding, Nutrition and Laboratory Animal Science | NA | Wrong study design |
| Estudio de los efectos de un modelo experimental sonoro y/o musical sobre la reactividad conductual y fisiológica de crías neonatales de rata Wistar | 2020 | Escribano et al. | Universidad Rey Juan Carlos | NA | Wrong exposure |
| Environmental enrichment: Music, day and dusk, how do they influence the rat's behaviour in laboratory (Rattus norvegicus)? | 2000 | Lemercier | STAL. Sciences et techniques de l'animal de laboratoire | NA | Full-text unavailable |
| A két oktávval megemelt (rágcsálók hallásához igazított) Mozart-szonáta hatása a patkányok tanulási képességére | 2015 | Horváth | Szent István University | NA | Full-text unavailable/Not in target language |
| Effect of combination of natural sounds during infancy on anxiety-like behaviors in adult rats | 2021 | Wang et al. | Journal of Jilin University (Medicine Edition) | 10.13481/j.1671-587X.20210409 | Full-text unavailable/Not in target language |
| Effects of Mozart Sonata on the rats' learning and memory performance. | 2014 | Fekete et al. | Magyar Állatorvosok Lapja | NA | Not in target language |
| Music exposure enhances resistance to Salmonella infection by promoting healthy gut microbiota | 2025 | Zhu et al. | Microbiology Spectrum | 10.1128/spectrum.02377-24 | Data not extractable |
| Music-Based Intervention Ameliorates Mecp2-Loss-Mediated Sociability Repression in Mice through the Prefrontal Cortex FNDC5/BDNF Pathway | 2021 | Hung et al. | International Journal of Molecular Sciences | 10.3390/ijms22137174 | Data not extractable |

| Title | Year | Authors | Journal | DOI | Reason for exclusion |
|---|---|---|---|---|---|
| Methamphetamine toxicity in mice is potentiated by exposure to loud music | 2001 | Morton et al. | Neuroreport | 10.1097/00001756-200110290-00026 | Data not extractable |
| Antidepressant effect of bright white LED combined with classical music | 2018 | Wang et al. | Infrared and Laser Engineering | 10.3788/IRLA201847.0420002 | Full text not available/Not in target language |
| Anxiolytic effect of Mozart music over short and long photoperiods as part of environmental enrichment in captive Rattus norvegicus (Rodentia: Muridae) | 2015 | Cruz et al. | Scandinavian Journal of Laboratory Animal Science | 10.23675/SJLAS.V41I0.341 | Data not extractable |
| Behavioral responses to emotional challenges in female rats living in a seminatural environment: The role of estrogen receptors | 2018 | Le Moëne and Ågmo | Hormones and Behavior | 10.1016/J.YHBEH.2018.10.013 | Wrong population |
| Critical period for acoustic preference in mice | 2012 | Yang et al. | Proceedings of the National Academy of Sciences of the United States of America | 10.1073/pnas.1200705109 | Wrong outcome/wrong study design |

### Appendix: PRISMA EcoEvo Checklist

O'Dea, R.E., Lagisz, M., Jennions, M.D., Koricheva, J., Noble, D.W., Parker, T.H., Gurevitch, J., Page, M.J., Stewart, G., Moher, D. and Nakagawa, S. (2021), Preferred reporting items for systematic reviews and meta-analyses in ecology and evolutionary biology: a PRISMA extension. Biol Rev. doi:10.1111/brv.12721

| Checklist item | Sub-item number | Sub-item | Reported by authors? | Notes |
|---|---|---|---|---|
| Title and abstract | 1.1 | Identify the review as a systematic review, meta-analysis, or both | Yes | |
| | 1.2 | Summarise the aims and scope of the review | Yes | |
| | 1.3 | Describe the data set | Yes | |
| | 1.4 | State the results of the primary outcome | Yes | |
| | 1.5 | State conclusions | Yes | |
| | 1.6 | State limitations | Yes | |
| Aims and questions | 2.1 | Provide a rationale for the review | Yes | Introduction |
| | 2.2 | Reference any previous reviews or meta-analyses on the topic | Yes | Introduction |
| | 2.3 | State the aims and scope of the review (including its generality) | Yes | Introduction |
| | 2.4 | State the primary questions the review addresses (e.g. which moderators were tested) | Yes | Methods. Section: Data collection |
| | 2.5 | Describe whether effect sizes were derived from experimental and/or observational comparisons | Yes | Methods. Section: Eligibility criteria and screening. Table 1. |
| Review registration | 3.1 | Register review aims, hypotheses (if applicable), and methods in a time-stamped and publicly accessible archive and provide a link to the registration in the methods section of the manuscript. Ideally registration occurs before the search, but it can be done at any stage before data analysis. | Yes | Methods. Section: Literature searching |
| | 3.2 | Describe deviations from the registered aims and methods | Yes | Methods: Section: Deviations from the preregistered protocol |
| | 3.3 | Justify deviations from the registered aims and methods | Yes | Methods: Section: Deviations from the preregistered protocol |
| Eligibility criteria | 4.1 | Report the specific criteria used for including or excluding studies when screening titles and/or abstracts, and full texts, according to the aims of the systematic review (e.g. study design, taxa, data availability) | Yes | Methods. Section: Eligibility criteria and screening |
| | 4.2 | Justify criteria, if necessary (i.e. not obvious from aims and scope) | NA | |

#### Appendix: PRISMA EcoEvo Checklist

| Checklist item | Sub-item number | Sub-item | Reported by authors? | Notes |
|---|---|---|---|---|
| Finding studies | 5.1 | Define the type of search (e.g. comprehensive search, representative sample) | Yes | Methods. Section: Literature searching |
| | 5.2 | State what sources of information were sought (e.g. published and unpublished studies, personal communications) | Yes | Methods. Section: Literature searching |
| | 5.3 | Include, for each database searched, the exact search strings used, with keyword combinations and Boolean operators | Yes | Supplementary material. Section: Literature searches |
| | 5.4 | Provide enough information to repeat the equivalent search (if possible), including the timespan covered (start and end dates) | Yes | Supplementary material. Section: Literature searches |
| Study selection | 6.1 | Describe how studies were selected for inclusion at each stage of the screening process (e.g. use of decision trees, screening software) | Yes | Methods. Section: Eligibility criteria and screening Preregistered protocol |
| | 6.2 | Report the number of people involved and how they contributed (e.g. independent parallel screening) | Yes | Methods. Section: Eligibility criteria and screening |
| Data collection process | 7.1 | Describe where in the reports data were collected from (e.g. text or figures) | Yes | Supplementary dataset |
| | 7.2 | Describe how data were collected (e.g. software used to digitize figures, external data sources) | Yes | Methods. Section: Data collection |
| | 7.3 | Describe moderator variables that were constructed from collected data (e.g. number of generations calculated from years and average generation time) | Yes | Methods: Section: Deviations from the preregistered protocol. Supplementary material table S1 |
| | 7.4 | Report how missing or ambiguous information was dealt with during data collection (e.g. authors of original studies were contacted for missing descriptive statistics, and/or effect sizes were calculated from test statistics) | Yes | Methods. Section: Data collection |
| | 7.5 | Report who collected data | Yes | Methods. Section: Data collection |
| | 7.6 | State the number of extractions that were checked for accuracy by co-authors | Yes | Methods. Section: Data collection |

### Appendix: PRISMA EcoEvo Checklist

| Checklist item | Sub-item number | Sub-item | Reported by authors? | Notes |
|---|---|---|---|---|
| Data items | 8.1 | Describe the key data sought from each study | Yes | Methods. Section: Data collection |
| | 8.2 | Describe items that do not appear in the main results, or which could not be extracted due to insufficient information | Yes | Methods. Section: Data collection<br>Methods: Section: Deviations from the preregistered protocol |
| | 8.3 | Describe main assumptions or simplifications that were made (e.g. categorising both 'length' and 'mass' as 'morphology') | Yes | Methods: Section: Deviations from the preregistered protocol |
| | 8.4 | Describe the type of replication unit (e.g. individuals, broods, study sites) | Yes | |
| Assessment of individual study quality | 9.1 | Describe whether the quality of studies included in the systematic review or meta-analysis was assessed (e.g. blinded data collection, reporting quality, experimental *versus* observational) | Yes | Methods. Section: Meta-analytic models |
| | 9.2 | Describe how information about study quality was incorporated into analyses (e.g. meta-regression and/or sensitivity analysis) | Yes | Methods. Section: Small-study effects, time-lag patterns, and sensitivity analyzes |
| Effect size measures | 10.1 | Describe effect size(s) used | Yes | Methods. Section: Effect size calculation |
| | 10.2 | Provide a reference to the equation of each calculated effect size (e.g. standardised mean difference, log response ratio) and (if applicable) its sampling variance | Yes | Methods. Section: Effect size calculation |
| | 10.3 | If no reference exists, derive the equations for each effect size and state the assumed sampling distribution(s) | Yes | Methods. Section: Effect size calculation |
| Missing data | 11.1 | Describe any steps taken to deal with missing data during analysis (e.g. imputation, complete case, subset analysis) | Yes | Methods. Section: Data collection<br>Online supplementary. Section: Effect size calculation |
| | 11.2 | Justify the decisions made to deal with missing data | Yes | Methods: Section: Deviations from the preregistered protocol |
| Meta-analytic | 12.1 | Describe the models used for synthesis of effect sizes | Yes | Methods. Section: Meta-analytic models |

| | | |
|---|---|---|
| model description | 12.2 | The most common approach in ecology and evolution will be a random-effects model, often with a hierarchical/multilevel structure. If other types of models are chosen (e.g. common/fixed effects model, unweighted model), provide justification for this choice |

| Checklist item | Sub-item number | Sub-item | Reported by authors? | Notes |
|---|---|---|---|---|
| Software | 13.1 | Describe the statistical platform used for inference (e.g. *R*) | Yes | Methods. Section: Statistical computing environment and software |
| | 13.2 | Describe the packages used to run models | Yes | Methods. Section: Statistical computing environment and software |
| | 13.3 | Describe the functions used to run models | Yes | Methods. Section: Statistical computing environment and software |
| | 13.4 | Describe any arguments that differed from the default settings | NA | |
| | 13.5 | Describe the version numbers of all software used | Yes | Methods. Section: Statistical computing environment and software |
| Non-independence | 14.1 | Describe the types of non-independence encountered (e.g. phylogenetic, spatial, multiple measurements over time) | Yes | Methods. Section: Meta-analytic models |
| | 14.2 | Describe how non-independence has been handled | Yes | Methods. Section: Meta-analytic models |
| | 14.3 | Justify decisions made | Yes | Methods. Section: Meta-analytic models |
| Meta-regression and model selection | 15.1 | Provide a rationale for the inclusion of moderators (covariates) that were evaluated in meta-regression models | Yes | Methods. Section: Data collection |
| | 15.2 | Justify the number of parameters estimated in models, in relation to the number of effect sizes and studies (e.g. interaction terms were not included due to insufficient sample sizes) | NA | |
| | 15.3 | Describe any process of model selection | NA | |
| Publication bias and sensitivity analyses | 16.1 | Describe assessments of the risk of bias due to missing results (e.g. publication, time-lag, and taxonomic biases) | Yes | Results. Section: Risk of bias and reporting transparency |
| | 16.2 | Describe any steps taken to investigate the effects of such biases (if present) | NA | |

| | | | | |
|---|---|---|---|---|
| | 16.3 | Describe any other analyses of robustness of the results, e.g. due to effect size choice, weighting or analytical model assumptions, inclusion or exclusion of subsets of the data, or the inclusion of alternative moderator variables in meta-regressions | Yes | Methods. Section: Effect size calculation |
| Clarification of *post hoc* analyses | 17.1 | When hypotheses were formulated after data analysis, this should be acknowledged. | | |

| **Checklist item** | **Sub-item number** | **Sub-item** | **Reported by authors?** | **Notes** |
|---|---|---|---|---|
| Metadata, data, and code | 18.1 | Share metadata (i.e. data descriptions) | Yes | Online supplementary material<br>GitHub repository |
| | 18.2 | Share data required to reproduce the results presented in the manuscript | Yes | GitHub repository |
| | 18.3 | Share additional data, including information that was not presented in the manuscript (e.g. raw data used to calculate effect sizes, descriptions of where data were located in papers) | Yes | Online supplementary material<br>GitHub repository |
| | 18.4 | Share analysis scripts (or, if a software package with graphical user interface (GUI) was used, then describe full model specification and fully specify choices) | Yes | Online supplementary material<br>GitHub repository |
| Results of study selection process | 19.1 | Report the number of studies screened | Yes | Supplementary material: flowchart |
| | 19.2 | Report the number of studies excluded at each stage of screening | Yes | Supplementary material: flowchart |
| | 19.3 | Report brief reasons for exclusion from the full text stage | Yes | Supplementary material table S4 |
| | 19.4 | Present a Preferred Reporting Items for Systematic Reviews and Meta-Analyses (PRISMA)-like flowchart (www.prisma-statement.org). | Yes | Supplementary material: flowchart |
| Sample sizes and study characteristics | 20.1 | Report the number of studies and effect sizes for data included in meta-analyses | Yes | Results. Section: Data overview. Figure 1. |
| | 20.2 | Report the number of studies and effect sizes for subsets of data included in meta-regressions | Yes | Results<br>Online supplementary material |

| | | | | |
|---|---|---|---|---|
| | 20.3 | Provide a summary of key characteristics for reported outcomes (either in text or figures; e.g. one quarter of effect sizes reported for vertebrates and the rest invertebrates) | Yes | Results. Section: Data overview. Figure 1. |
| | 20.4 | Provide a summary of limitations of included moderators (e.g. collinearity and overlap between moderators) | NA | |
| | 20.5 | Provide a summary of characteristics related to individual study quality (risk of bias) | Yes | Results.Section: Risk of bias and reporting transparency |

| Checklist item | Sub-item number | Sub-item | Reported by authors? | Notes |
|---|---|---|---|---|
| Meta-analysis | 21.1 | Provide a quantitative synthesis of results across studies, including estimates for the mean effect size, with confidence/credible intervals | Yes | Results. Section: Overall effects of music exposure |
| Heterogeneity | 22.1 | Report indicators of heterogeneity in the estimated effect (e.g. $I^2$, $tau^2$ and other variance components) | Yes | Results. |
| Meta-regression | 23.1 | Provide estimates of meta-regression slopes (i.e. regression coefficients) and confidence/credible intervals | Yes | Results |
| | 23.2 | Include estimates and confidence/credible intervals for all moderator variables that were assessed (i.e. complete reporting) | Yes | Results<br>Online supplementary material |
| | 23.3 | Report interactions, if they were included | | |
| | 23.4 | Describe outcomes from model selection, if done (e.g. R2 and AIC) | | |
| Outcomes of publication bias and sensitivity analyses | 24.1 | Provide results for the assessments of the risks of bias (e.g. Egger's regression, funnel plots) | Yes | Results. Section: Risk of bias and reporting transparency |
| | 24.2 | Provide results for the robustness of the review's results (e.g. subgroup analyses, meta-regression of study quality, results from alternative methods of analysis, and temporal trends) | Yes | Supplementary material Figure S3.<br>Online supplementary material |
| Discussion | 25.1 | Summarise the main findings in terms of the magnitude of effect | Yes | Results |

| | | | | |
|---|---|---|---|---|
| | 25.2 | Summarise the main findings in terms of the precision of effects (e.g. size of confidence intervals, statistical significance) | | |
| | 25.3 | Summarise the main findings in terms of their heterogeneity | Yes | Results |
| | 25.4 | Summarise the main findings in terms of their biological/practical relevance | Yes | Results. Table 2 |
| | 25.5 | Compare results with previous reviews on the topic, if available | Yes | Discussion. Section: Music exposure in the context of environmental enrichment |
| | 25.6 | Consider limitations and their influence on the generality of conclusions, such as gaps in the available evidence (e.g. taxonomic and geographical research biases) | Yes | Section: Study-specific limitations and recommendations for future research |

| Checklist item | Sub-item number | Sub-item | Reported by authors? | Notes |
|---|---|---|---|---|
| Contributions and funding | 26.1 | Provide names, affiliations, and funding sources of all co-authors | Yes | |
| | 26.2 | List the contributions of each co-author | Yes | |
| | 26.3 | Provide contact details for the corresponding author | Yes | |
| | 26.4 | Disclose any conflicts of interest | Yes | |
| References | 27.1 | Provide a reference list of all studies included in the systematic review or meta-analysis | Yes | Main references. Supplementary material table S3 |
| | 27.2 | List included studies as referenced sources (e.g. rather than listing them in a table or supplement) | Yes | Main references |

### References

Tingting An, Zhenhua Song, and Jin-hui Wang. Molecular mechanism of reward treatment ameliorating chronic stress-induced depressive-like behavior assessed by sequencing miRNA and mRNA in medial prefrontal cortex. *Biochemical and Biophysical Research Communications*, 528(3):520–527, 7 2020. ISSN 0006291X. doi: 10.1016/j.bbrc.2020.05.158. URL https://linkinghub.elsevier.com/retrieve/pii/S0006291X20310962.

Anice Milbratz de Camargo, Daniela Delwing de Lima, Débora Delwing Dal Magro, Johanna Kleis Seubert, Júlia Niehues da Cruz, and José Geraldo Pereira da Cruz. Adjuvant effects of classical music on simvastatin induced reduction of anxiety but not object recognition memory in rats. *Psychology & Neuroscience*, 6(3):403–410, 7 2013. ISSN 1983-3288. doi: 10.3922/j.psns.2013.3.19. URL https://doi.apa.org/doi/10.3922/j.psns.2013.3.19.

Si Chen, Tuo Liang, Fiona H. Zhou, Ye Cao, Chao Wang, Fei-Yifan Wang, Fang Li, Xin-Fu Zhou, Jian-Yi Zhang, and Chang-Qi Li. Regular Music Exposure in Juvenile Rats Facilitates Conditioned Fear Extinction and Reduces Anxiety after Foot Shock in Adulthood. *BioMed Research International*, 2019:1–10, 7 2019. ISSN 2314-6133. doi: 10.1155/2019/8740674. URL https://www.hindawi.com/journals/bmri/2019/8740674/.

Hong-Yu Cheng, Hao-Xue Xie, Qian-Lan Tang, Li-Tao Yi, and Ji-Xiao Zhu. Light and classical music therapies attenuate chronic unpredictable mild stress-induced depression via BDNF signaling pathway in mice. *Heliyon*, 10(13):e34196, 7 2024. ISSN 24058440. doi: 10.1016/j.heliyon.2024.e34196. URL https://linkinghub.elsevier.com/retrieve/pii/S2405844024102277.

Sachiko Chikahisa, Atsuko Sano, Kazuyoshi Kitaoka, Ken-Ichi Miyamoto, and Hiroyoshi Sei. Anxiolytic effect of music depends on ovarian steroid in female mice. *Behavioural Brain Research*, 179(1):50–59, 4 2007. ISSN 01664328. doi: 10.1016/j.bbr.2007.01.010. URL https://linkinghub.elsevier.com/retrieve/pii/S0166432807000289.

J N Cruz, D D Lima, D D Dal Margo, and J G P Cruz. Anxiolytic effect of Mozart music over short and long photoperiods as part of environmental enrichment in captive Rattus norvegicus (Rodentia: Muridae). *Scandinavian Journal of Laboratory Animal Science*, 41:1–7, 2015. ISSN 2002-0112. doi: 10.23675/SJLAS.V41I0.341. URL https://ojs.utlib.ee/index.php/SJLAS/article/view/21615/16318.

Begoña Escribano, Ismael Quero, Montserrat Feijóo, Inmaculada Tasset, Pedro Montilla, and Isaac Túnez. Role of noise and music as anxiety modulators: Relationship with ovarian hormones in the rat. *Applied Animal Behaviour Science*, 152:73–82, 3 2014. ISSN 01681591. doi: 10.1016/j.applanim.2013.12.006. URL https://linkinghub.elsevier.com/retrieve/pii/S0168159113002955.

Emilio Mateu Escribano, Agustín Martínez Peláez, Manuel Sánchez Cid, and José Antonio Martínez Orgado. *Estudio de los efectos de un modelo experimental sonoro y/o musical sobre la reactividad conductual y fisiológica de crías neonatales de rata Wistar*. PhD thesis, Universidad Rey Juan Carlos, 2020.

Oliver Falkenhorst. *Is Mice Behavior Influenced by Music?* PhD thesis, Szent István University, 2013.

S. Fekete and T. Bernitsa. Influence of different environmental effects of human origin (socialization, music, noisemusic, noise) upon the rats' behaviour. Part 2. Do rats react on human music? *Magyar Állatorvosok Lapja*, 135(4), 4 2013.

S. Fekete, Chihiro Sukikara, and G. Korsós. Influence of different environmental effects of human origin (socialization, music, noisemusic, noise) upon the rats' behaviour. Part 3. Effect of different noises on the open-field test behaviour. *Magyar Állatorvosok Lapja*, 135(11), 11 2013.

S. Fekete, A. Lukács, K. Horváth, G. Korsós, and T. Vezér. Effects of Mozart Sonata on the rats' learning and memory performance. *Magyar Állatorvosok Lapja*, 136(3), 2014. URL `http://hdl.handle.net/10832/1445`.

Montse Flores-García, África Flores, Ester Aso, Paloma Otero-López, Francisco Ciruela, Sebastià Videla, Jennifer Grau-Sánchez, Antoni Rodríguez-Fornells, Jordi Bonaventura, and Víctor Fernández-Dueñas. Dopamine dynamics in chronic pain: music-induced, sex-dependent, behavioral effects in mice. *Pain Reports*, 10(1):e1205, 2 2024. doi: 10.1097/PR9.0000000000001205. URL `https://pmc.ncbi.nlm.nih.gov/articles/PMC11631031/`.

Enrique Flores-Gutiérrez, Edith Araceli Cabrera-Muñoz, Nelly Maritza Vega-Rivera, Leonardo Ortiz-López, and Gerardo Bernabé Ramírez-Rodríguez. Exposure to Patterned Auditory Stimuli during Acute Stress Prevents Despair-Like Behavior in Adult Mice That Were Previously Housed in an Enriched Environment in Combination with Auditory Stimuli. *Neural Plasticity*, 2018:1–14, 12 2018. ISSN 2090-5904. doi: 10.1155/2018/8205245. URL `https://www.hindawi.com/journals/np/2018/8205245/`.

Breno Raul Freitas Oliveira, Tiago Werley Pires da Silva, Raissa Maria Carvalho Alves, José Ribamar Soares Neto, Camila Araújo Oliveira, Daniela Feliz Barbosa Guerreiro, Mariana Mazza Santos, Emanuel Ramos da Costa, Cristovam Wanderley Picanço Diniz, and Daniel Guerreiro Diniz. Classical Music and Environmental Enrichment Enhanced Spatial Memory and Learning and Increased Mouse Innate Tendency to Avoid Open Spaces. *EC Neurology*, 13:12–20, 2020. doi: 10.31080/ecne.2021.13.00837.

Qiang Fu, Rui Qiu, Lei Chen, Yuewen Chen, Wen Qi, and Yong Cheng. Music prevents stress-induced depression and anxiety-like behavior in mice. *Translational Psychiatry*, 13(1):317, 10 2023. ISSN 2158-3188. doi: 10.1038/s41398-023-02606-z. URL `https://www.nature.com/articles/s41398-023-02606-z`.

Qiang Fu, Rui Qiu, Tongtong Yao, Liming Liu, Yaobo Li, Xiaodong Li, Wen Qi, Yuewen Chen, and Yong Cheng. Music therapy as a preventive intervention for postpartum depression: modulation of synaptic plasticity, oxidative stress, and inflammation in a mouse model. *Translational Psychiatry*, 15(1):143, 4 2025. ISSN 2158-3188. doi: 10.1038/s41398-025-03370-y. URL `https://www.nature.com/articles/s41398-025-03370-y`.

Elena Alexandrovna Germanovich. Vliyanie razlichnykh zvukovykh stressorov na povedenie laboratornykh myshei v teste ”otkrytoe pole”. *Aktual'nye problemy biologicheskoi i khimicheskoi ekologii*, pages 146–149, 2016.

K. Horváth. *A két oktávval megemelt (rágcsálók hallásához igazított) Mozart-szonáta hatása a patkányok tanulási képességére*. PhD thesis, Szent István University, 2015.

Pi-Lien Hung, Kay L. H. Wu, Chih-Jen Chen, Ka-Kit Siu, Yi-Jung Hsin, Liang-Jen Wang, and Feng-Sheng Wang. Music-Based Intervention Ameliorates Mecp2-Loss-Mediated Sociability Repression in Mice through the Prefrontal Cortex FNDC5/BDNF Pathway. *International Journal of Molecular Sciences*, 22(13):7174, 7 2021. ISSN 1422-0067. doi: 10.3390/ijms22137174. URL `https://www.mdpi.com/1422-0067/22/13/7174`.

Qian Ji, Zhi Gang Li, Yin Shan Tang, Yu Ping Mo, Hai Jiang Yao, and Chao Ketu Saiyin. [Effect of electroacupuncture intervention on behavioral changes and hippocampal excitatory amino acid transporter mRNA expression in depression rats]. *Zhen Ci Yan Jiu*, 38(3), 2013. ISSN 1000-0607. URL `https://pubmed.ncbi.nlm.nih.gov/24006665/`.

G. Korsós and S. Fekete. Influence of different environmental effects of human origin (socialization, music, noisemusic, noise) upon the rats' behaviour. Part 1. *Magyar Állatorvosok Lapja*, 135(2), 2013.

G. Korsós, D. Brown, C. Windig-Zavadil, S. Ruhlickle, and S. Fekete. Mouse Ethology: Effect of different human and rodentized music upon the social and individual behaviour, general feeling and genetics-environment interaction of mice II. Effect of original and five octaves higher music of Bach and MOZART on the behaviour of mice of different genotypes. *Magyar Állatorvosok Lapja*, 138(12), 2016.

Tanuja Krishnamurthy and Bhagya Venkanna Rao. Indian classical Mohana Raga (instrumental music) overcomes anxiety, depression and memory impairment in chronic unpredictable mild stress rat model –A behavioural study. *Indian Journal of Traditional Knowledge*, 4 2025. ISSN 09725938. doi: 10.56042/ijtk.v24i4.12620. URL `https://or.niscpr.res.in/index.php/IJTK/issue/view/386`.

Shelly Kusumarini, Lita Rakhma Yustinasari, Eka Pramystha Hestianah, Suryo Kuncorojati, and Tutik Juniastuti. Mozart KV 448 Menurunkan Densitas dan Aktivitas Neuroglia Hipokampus Mencit (Mus musculus) Selama Stres Prenatal No. 416-KE. *Jurnal Sain Veteriner*, 35(1):1, 10 2017. ISSN 2407-3733. doi: 10.22146/jsv.29279. URL `https://jurnal.ugm.ac.id/jsv/article/view/29279`.

Olivia Le Moëne and Anders Ågmo. Behavioral responses to emotional challenges in female rats living in a seminatural environment: The role of estrogen receptors. *Hormones and Behavior*, 106:162–177, 11 2018. ISSN 0018-506X. doi: 10.1016/J.YHBEH.2018.10.013. URL `https://www.sciencedirect.com/science/article/`

abs/pii/S0018506X1830206X.

H. Lemercier. Environmental enrichment: Music, day and dusk, how do they influence the rat's behaviour in laboratory (Rattus norvegicus)? *STAL. Sciences et techniques de l'animal de laboratoire*, 25(2), 1 2000.

Wen Jing Li, Hui Yu, Jian Min Yang, Jing Gao, Hong Jiang, Min Feng, Yu Xia Zhao, and Zhe Yu Chen. Anxiolytic effect of music exposure on BDNFMet/Met transgenic mice. *Brain Research*, 1347:71–79, 8 2010. ISSN 00068993. doi: 10.1016/j.brainres.2010.05.080.

Tuo Liang, Chao Wang, Ye Cao, Feiyifan Wang, Si Cheb, Mei Zheng, and Changqi Li. The effect of music exposure in juvenile stage on anxiety-like behavior and fear extinction in adult rat. *Chinese Journal of Behavioral Medicine and Brain Science*, pages 311–314, 2012. doi: 10.3760/CMA.J.ISSN.1674-6554.2012.04.008. URL http://dx.doi.org/10.3760/cma.j.issn.1674-6554.2012.04.008.

F. Lin, Y. Wu, B. Yuan, W. Feng, and Y. Bian. Musicotherapy treating rat models with post-stroke depression by reducing expressions of serum IL-10 and hippocampal PDE4A. *BASIC & CLINICAL PHARMACOLOGY & TOXICOLOGY*, 125, 2019.

Q. Lopez, F. Tunez, and P. Montilla. ROLE OF HO-1 IN THE EFFECTOR PHASE OF ARTHRITIS FINASTERIDE AND ALLO-PREGENENOLONE ADMINISTRATION ON THE ANXIOLYTIC EFFECT OF MUSIC IN FEMALE RATS. INTRODUCTION. *BASIC & CLINICAL PHARMACOLOGY & TOXICOLOGY*, 109, 2011.

Anice Milbratz de Camargo, Henrique Bonde, Débora Delwing Dal Magro, Daniela Delwing de Lima, and Luciane Coutinho de Azevedo Campanella. Cocoa and classical music: effect on anxiety and antioxidant activity in Wistar rats. *Archivos Latinoamericanos de Nutrición*, 67(2), 2017.

D. Monje-Reyna, L García-Hernández, G. Coria-Ávila, M. Toledo, M. Hernández-Aguilar, and J. Manzo-Denes. Patterns of movement in the open field of the rat with autistic behavior subjected to musical stimulation. *Neurobiología*, 9 (20), 2018.

A. Jennifer Morton, Miriam A. Hickey, and Laura C. Dean. Methamphetamine toxicity in mice is potentiated by exposure to loud music. *Neuroreport*, 12(15):3277–3281, 10 2001. ISSN 0959-4965. doi: 10.1097/00001756-200110290-00026. URL http://journals.lww.com/00001756-200110290-00026.

Júlia Niehues da Cruz, Daniela Delwing de Lima, Débora Delwing Dal Magro, and José Geraldo Pereira da Cruz. The Power of Classic Music to Reduce Anxiety in Rats Treated with Simvastatin. *Basic and Clinical Neuroscience*, 2(4): 5–11, 2011. URL https://bcn.iums.ac.ir/article-1-173-en.html.

Lisa Pangemanan, Irwanto, and Margarita M. Maramis. Mozart K488 Addition Can Improve Depressive-Like Behavior in Rats: In Search of Better Management. *Pharmacognosy Journal*, 16(2):348–354, 3 2024. ISSN 09753575. doi:

10.5530/pj.2024.16.53.

A. Papadakakis, K. Sidiropoulou, and G. Panagis. Music exposure attenuates anxiety- and depression-like behaviors and increases hippocampal spine density in male rats. *Behavioural Brain Research*, 372:112023, 10 2019. ISSN 01664328. doi: 10.1016/j.bbr.2019.112023. URL `https://linkinghub.elsevier.com/retrieve/pii/S0166432819303249`.

Ann M. Peiffer, Glenn D. Rosen, and R.Holly Fitch. Rapid auditory processing and MGN morphology in microgyric rats reared in varied acoustic environments. *Developmental Brain Research*, 138(2):187–193, 10 2002. ISSN 01653806. doi: 10.1016/S0165-3806(02)00472-8. URL `https://linkinghub.elsevier.com/retrieve/pii/S0165380602004728`.

J.E. Polston, H.Y. Rubbinaccio, J.T. Morra, E.M. Sell, and S.D. Glick. Music and methamphetamine: Conditioned cue-induced increases in locomotor activity and dopamine release in rats. *Pharmacology Biochemistry and Behavior*, 98 (1):54–61, 3 2011. ISSN 00913057. doi: 10.1016/j.pbb.2010.11.024. URL `https://linkinghub.elsevier.com/retrieve/pii/S009130571000362X`.

Pratiksha Rathod and Harsha D Vaidya. Study of the effect of Kalyani raga in Anxiety-like conditions in female Wistar rats. *International Journal of Ayurvedic Medicine*, 15(1):30–34, 4 2024. ISSN 0976-5921. doi: 10.47552/ijam.v15i1.2977. URL `https://ijam.co.in/index.php/ijam/article/view/2977`.

Jingyao Ren and Jian Lu. Heavy metal Music, Hip-hop Music and Construction Noise Induces Depressive Symptoms in mice. *ASEAN Journal of Psychiatry*, 25(8):1–14, 10 2024.

Lou Rizzolo, Marianne Leger, Sophie Corvaisier, Mathilde Groussard, Hervé Platel, Valentine Bouet, Pascale Schumann-Bard, and Thomas Freret. Long-Term Music Exposure Prevents Age-Related Cognitive Deficits in Rats Independently of Hippocampal Neurogenesis. *Cerebral Cortex*, 31(1):620–634, 1 2021. ISSN 1047-3211. doi: 10.1093/cercor/bhaa247. URL `https://academic.oup.com/cercor/article/31/1/620/5909651`.

Hasan Saghari, Vahid Sheibani, Khadijeh Esmaeilpour, and Naeem ur Rehman. Music Alleviates Learning and Memory Impairments in an Animal Model of Post-Traumatic Stress Disorder. *Biointerface Research in Applied Chemistry*, 11(1):7775–7784, 7 2021. ISSN 2069-5837. doi: 10.33263/BRIAC111.77757784. URL `https://biointerfaceresearch.com/wp-content/uploads/2020/07/20695837111.77757784.pdf`.

Waneli Cristine Morais Sampaio, Mara Cláudia Ribeiro, Larice Feitosa Costa, Wânia Cristina de Souza, Goiara Mendonça de Castilho, Melissa Sousa de Assis, Fabiana Pirani Carneiro, Domitilla Marchiori, Nadyelle Targino de Lima, Pacífica Pinheiro Cavalcanti, and Vania Moraes Ferreira. Effect of music therapy on the developing central nervous system of rats. *Psychology & Neuroscience*, 10(2):176–188, 6 2017. ISSN 1983-3288. doi: 10.1037/pne0000087.

URL https://doi.apa.org/doi/10.1037/pne0000087.

Chihiro Sukikara. *How the Different Noise Types May Influence the Open-field Behavior of Rats?* PhD thesis, Szent István University, 2013. URL http://hdl.handle.net/10832/943.

Gamze Tümentemur, Mustafa Titiz, Sümeyye Çilingir, Gülşah Ertosun, Dilan Acar, Mehmet Yavuz, and Guldal Süyen. Investigation of the Influence of Music on Behavioral Findings by Valproic Acid-Induced Autism Model in Rats. In *Turkish Society of Physiological Sciences 44th National Physiology Congress*, volume 225, 2018.

Şule Terzioglu Uşak, Aleyna Dal, Hilal Yanik, and Birsen Elibol. Effects of music on stress induced hormones and oxidative stress levels. *Cukurova Medical Journal*, 45(4):1493–1498, 12 2020. ISSN 2602-3032. doi: 10.17826/cumj.735738. URL http://dergipark.org.tr/en/doi/10.17826/cumj.735738.

Danni Wang, Yuhuan Dai, Jing Cheng, Dawei Zhang, Cheng Wang, and Songlin Zhuang. Antidepressant effect of bright white LED combined with classical music. *Infrared and Laser Engineering*, 47(4):0420002, 4 2018. ISSN 1007-2276. doi: 10.3788/IRLA201847.0420002. URL https://www.researching.cn/articles/OJef5287faf6fef1a5/html.

Qian Wang, Siyue Zhuang, Guilong Wu, and Shengtian Li. Effect of combination of natural sounds during infancy on anxiety-like behaviors in adult rats. *Journal of Jilin University (Medicine Edition)*, 47(4), 2021. doi: 10.13481/j.1671-587X.20210409.

Eun Jin Yang, Eric W. Lin, and Takao K. Hensch. Critical period for acoustic preference in mice. *Proceedings of the National Academy of Sciences of the United States of America*, 109(SUPPL.2):17213–17220, 10 2012. ISSN 00278424. doi: 10.1073/pnas.1200705109.

Clara Y. Zhu, Hyuntae Byun, Elyza A. Do, Yue Zhang, Ethan Tanchoco, Joris Beld, Ansel Hsiao, and Jun Zhu. Music exposure enhances resistance to Salmonella infection by promoting healthy gut microbiota. *Microbiology Spectrum*, 13(5), 5 2025. ISSN 2165-0497. doi: 10.1128/spectrum.02377-24. URL https://journals.asm.org/doi/10.1128/spectrum.02377-24.
